# Cell splitting in *Staphylococcus aureus* is controlled by an adaptor protein facilitating degradation of a peptidoglycan hydrolase

**DOI:** 10.1101/2025.05.11.653310

**Authors:** Maria Disen Barbuti, Eivind Frøyland Skjennum, Viktor Hundtofte Mebus, Danae Morales Angeles, Matilde Hustad, Marita Torrissen Mårli, Dorte Frees, Morten Kjos

## Abstract

Regulated protein degradation by Clp proteases is a highly conserved post-translational control mechanism in bacteria. In *Staphylococcus aureus*, the ClpXP complex targets the peptidoglycan hydrolase Sle1, maintaining a tightly regulated balance between peptidoglycan biosynthesis and hydrolysis, which is required to ensure proper cell splitting without compromising cell integrity. β-lactams antibiotics disturb this balance, leading to their bactericidal effects. The mechanism underlying the specific targeting of Sle1 by the conserved ClpXP complex remains unknown. From a genome-wide screen for determinants of penicillin G susceptibility in *S. aureus*, we here identify the uncharacterized protein CxaR (for <u>C</u>lp<u>X</u>P-associated <u>a</u>utolytic regulator). Growth defects, premature cell splitting, and increased cell lysis were observed in the absence of CxaR. Interestingly, these defects were mitigated by sublethal concentrations of β-lactams. Through sequencing *cxaR* suppressor mutants, followed by immunoblotting, we show that the *cxaR* phenotypes are caused by excessive Sle1 accumulation. Indeed, exposure to β-lactams reduces Sle1 levels, thereby rescuing the cells lacking CxaR. Furthermore, *in vivo* protein-protein interaction assays demonstrated that CxaR directly interacts with both ClpXP and Sle1, whereas no direct interaction was detected between Sle1 and ClpX. In line with this, CxaR was found to co-localize with ClpX adjacent to the septum. Taken together, these findings reveal that CxaR is a new regulatory factor controlling staphylococcal cell splitting by acting as an adaptor protein for controlled ClpXP-mediated degradation of Sle1.

## Introduction

Bacterial proliferation depends on the precise coordination of cell division, a complex process that culminates in the separation of daughter cells. In the Gram-positive coccoid pathogen *Staphylococcus aureus*, division initiates with the synthesis of a septal cross-wall at midcell, generating two hemispherical daughter cells that remain transiently connected via a peripheral wall ring [1–4]. Only after the septum is fully formed does this septum-adjacent peripheral cell wall undergo rapid resolution (“popping”) [1, 2]. This step, occurring in less than two milliseconds, is highly regulated to prevent premature splitting of the septum, which would compromise cell integrity and viability, ultimately leading to cell lysis [5, 6]. Septal splitting is facilitated by peptidoglycan hydrolases (PGHs) that create small perforations in the peripheral peptidoglycan ring, causing mechanical crack propagation [1, 7]. *S. aureus* encodes at least 20 PGHs with different physiological roles during bacterial growth and division [5, 8]. Among these, the amidase Sle1, which cleaves the amide bond linking the stem peptide to the glycan chain, plays a particularly important role in cell splitting, as its absence results in cell clustering, delays in cell cycle progression, and the formation of tetrad cells [2, 7, 9, 10]. Sle1 is translocated to the extracellular milieu by active secretion, facilitated by its N-terminal signal peptide. Three cell wall-binding LysM domains ensure localization to the division septum, where the catalytic CHAP domain facilitates peptidoglycan cleavage [11, 12].

As with all PGHs, Sle1 activity must be tightly regulated both spatially and temporally to prevent unintended breaches in the cell wall [5, 6]. Targeted recruitment of these enzymes to specific regions of the cell serves as a key regulatory mechanism in many bacteria, including *S. aureus* [11, 13, 14]. The localization of Sle1 is dependent on the anionic glycopolymers wall teichoic acids (WTAs). The reduced levels of WTA at the septum promote Sle1 accumulation and peptidoglycan cleavage in this region [11, 15]. The two-component systems WalKR and GraSR, facilitates transcriptional regulation of Sle1, ensuring proper coordination of cell wall remodeling and division [16–19]. Furthermore, recent studies have identified the DNA translocase and cell division protein FtsK [10] as well as a non-coding RNA named Rbc1 [20] as positive regulators of Sle1. Finally, and importantly, Sle1 levels are regulated by the ClpXP protease through targeted degradation [7].

Regulated protein degradation by Clp proteases affects all biological pathways, playing a crucial role in both general protein quality control and the targeted degradation of specific proteins in bacterial cells [21–23]. The ClpXP complex consist of the ATP-dependent hexameric ClpX ring, which recognizes, unfolds, and translocates specific protein substrates into the ClpP proteolytic chamber for degradation [24]. While ClpXP can directly target various substrates, efficient recognition of certain proteins requires separately encoded adaptor proteins that recognize specific substrates and assist in their delivery to ClpXP for degradation [21–23]. The vital roles of ClpXP adaptor proteins have been characterized in different bacteria, including *Escherichia coli* with SspB [25–27], UmuD [28, 29], and RssB [30, 31]. The only known adaptor protein for ClpXP in *S. aureus* is YjbH, which targets the stress-associated transcriptional regulator Spx for degradation [32–34], similar to its role in *B. subtilis* [35, 36].

Only a few ClpXP substrates have been verified in *S. aureus*, including Spx and Sle1 [37–39]. Unlike Spx, the mechanism by which ClpXP recognizes Sle1 is unknown. Nonetheless, the targeted degradation of Sle1 has been shown to be crucial for the survival of staphylococcal cells [7, 39–41]. Deletion of *clpX* results in abnormal septum formation and premature splitting of daughter cells [40–42]. This phenotype is mainly attributed to the impaired proteolytic regulation and subsequent accumulation of PGHs, particularly Sle1 [7, 38–40, 42, 43]. Beyond proteolysis, ClpX also appears to contribute to cell division independently of ClpP, as only *clpX* mutants exhibit heat stability and cold sensitivity [39, 44], a temperature dependent growth defect that can be alleviated by a mutation in the essential cell division protein FtsA [43]. ClpX has additionally been found to impact transcription of genes involved in peptidoglycan synthesis and cell division independently of ClpP [45]. Interestingly, addition of β-lactams, inhibition of lipoteichoic acid (LTA) and wall teichoic acid (WTA) biosynthesis, or inactivation of Sle1 can rescue growth of *clpX* mutants [41, 42], further evidencing a tight interplay between ClpXP, Sle1 and cell wall biogenesis in *S. aureus*.

The wide range of regulatory mechanisms invested into governing PGHs, and particularly Sle1, underlines the importance of maintaining control of these enzymes. Indeed, the bactericidal mechanism of β-lactam antibiotics appears to involve the dysregulation of PGH activity [46]. β-lactams inhibit the peptidoglycan cross-linking capacity of penicillin-binding proteins (PBPs) by covalently binding their active site [47–49]. This is believed to disturb the delicate balance between polymerization and autolytic degradation of peptidoglycan, which ultimately result in cell lysis [46, 50–52]. The relationship between PBPs and PGHs appears to be highly complex. On one hand, low-dose treatment with β-lactams result in thickened septal walls, delay cell separation, and significantly reduce the levels of specific PGHs, including Sle1 [7, 41, 53–56]. These findings suggest that β-lactams may exert their bactericidal effects, in part, by lowering the activity of specific PGHs, resulting in aberrant septum formation that does not permit further cell growth and division. [49, 50, 57]. On the other hand, β-lactam treatment has frequently been associated with activation of PGHs, leading to excessive cell wall degradation and subsequent cell lysis [50–52, 57]. Studies in various bacteria, including *Streptococcus pneumoniae* and *Bacillus subtilis*, have shown that when autolytic activity is suppressed, exposure to penicillin primarily results in growth inhibition rather than lysis [49, 50, 57]. The contrasting observations underscore that the exact mode of action of β-lactams is more complex than initially assumed and may depend on species-specific factors.

In the current study, we show that SAOUHSC_00659, a hitherto uncharacterized protein here renamed CxaR (for <u>C</u>lp<u>X</u>P-associated <u>a</u>utolytic regulator), controls cellular levels of Sle1 by mediating its targeted degradation by the ClpXP complex. Deletion or knockdown of the gene encoding CxaR resulted in accumulation of Sle1, which in turn contributed to growth deficiencies, accelerated daughter cell splitting, and increased cell lysis. Co-localization analyses combined with protein-protein interaction assays revealed that CxaR depends on ClpX for its septal-adjacent localization. While both proteins directly interact with each other, our results suggest that only CxaR interacts with Sle1. Together, these findings identify and elucidate the mechanism of a new adaptor protein required for spatiotemporal regulation of bacterial cell splitting.

## Results

### Sublethal concentrations of cell wall targeting antibiotics stimulate growth of cells lacking *cxaR*

A genome-wide, pooled CRISPR interference sequencing (CRISPRi-seq) screen [58–60] was conducted to identify genetic factors modulating susceptibility to β-lactams in *S. aureus* NCTC8325-4 (see **S1 Table** and **S1 Fig**). By knocking down the expression of transcriptional units, both in the presence and absence of inhibitory concentrations of penicillin G, we could quantify the associated growth fitness cost for each unit. The most prominent hit from this screen was a gene of unknown function, *SAOUHSC_00659* (hereafter referred to as *cxaR*), whose knockdown led to reduced fitness, an effect that appeared to be counteracted by the presence of penicillin G (**S1 Fig**).

To validate this result, a *cxaR* CRISPRi knockdown strain was created in *S. aureus* NCTC8325-4. CxaR depletion resulted in a distinct growth profile in TSB at 37°C, characterized by reduced initial growth rate, followed by a transient decline in optical density (OD), and a subsequent gradual recovery of growth (**Fig 1A**). The growth defect of *cxaR* knockdown was similar, although even more pronounced, in the MRSA strain JE2 (**Fig 1A**). A *cxaR* deletion mutant was also constructed in NCTC8325-4 by allelic exchange with a spectinomycin resistance cassette. The resulting strain (NCTC8325-4 Δ*cxaR*::*spc*) exhibited the same biphasic growth (**Fig 1B**), confirming that the growth phenotype was not caused by the CRISPRi-system. Finally, to exclude the possibility that the phenotype observed upon knockdown or deletion of *cxaR* was caused by polar effect on neighboring genes, we designed sgRNAs for CRISPRi knockdown of five adjacent genetic loci, located both upstream and downstream of *cxaR* (**S2A Fig**). Growth analyses revealed that none of these strains exhibited growth patterns similar to the *cxaR* mutant (**S2B-C Fig**), confirming that the observed effects are specifically attributable to *cxaR*.

**Fig. 1.**
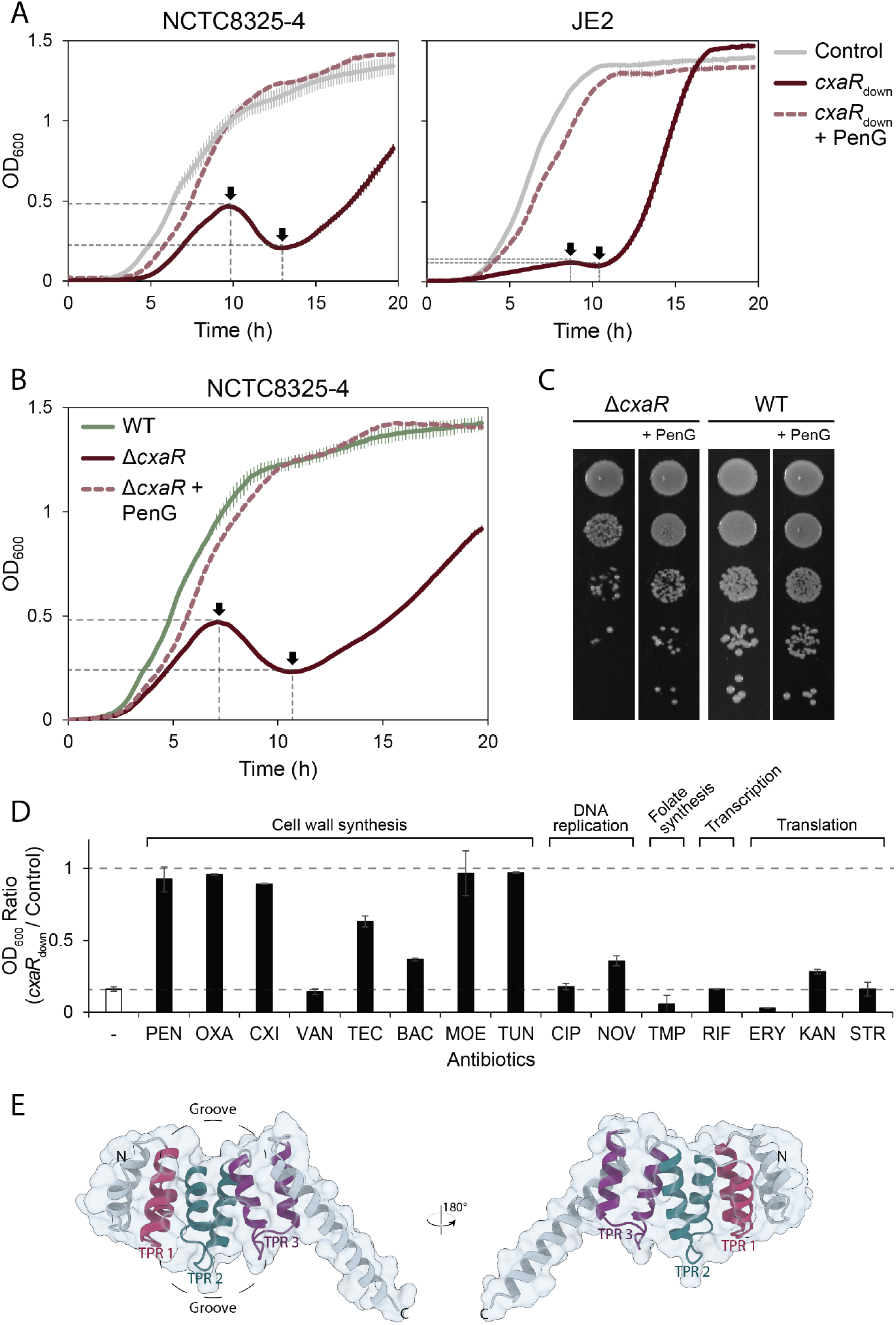
Lack of CxaR leads to a biphasic growth defect that can be rescued by sublethal concentrations of various antibiotics. (**A**) Growth curves of the *cxaR* knockdown mutants (*cxaR*_down_) and CRISPRi control strains in TSB at 37°C for both NCTC8325-4 and JE2. Gene knockdown was induced with addition of 500 μM IPTG, and growth of the *cxaR*_down_ mutants was restored with addition of penicillin G (1 ng/mL for NCTC8325-4 and 5 ng/mL for JE2). The graphs represent averages with standard deviation from triplicate measurements. (**B**) Growth curves of the NCTC8325-4 *cxaR* deletion mutant and wild-type in TSB at 37°C. Growth of the Δ*cxaR* mutant was restored with addition of penicillin G (1 ng/mL). The graphs represent averages with standard deviation from triplicate measurements. (**C**) Growth of the NCTC8325-4 *cxaR* deletion mutant and wild-type on TSA with or without penicillin G (1 ng/mL) after 10 hours of incubation at 37°C. The bacterial cultures were sampled after 10 hours of incubation in TSB (± penicillin G) and 10-fold dilution series were spotted onto TSA plates (± penicillin G). (**D**) Relative growth of the JE2 *cxaR*_down_ mutant compared to the CRISPRi control after 8 hours of incubation at 37°C in the presence or absence of MIC_1_ concentrations of various antibiotics. Data represent the average growth ratio ± standard deviation from the mean of two biological replicates for the antibiotic-treated samples, and from 30 biological replicates for the untreated control. A ratio of 1 indicates that the two strains have reached the exact same OD_600_ after 8 hours of incubation. In the absence of antibiotics, the ratio was approximately 0.16. Notably, several of the antibiotics stimulated growth of the *cxaR*_down_ mutant, leading to a significant increase in the growth ratio. For the individual growth of the *cxaR*_down_ and CRISPRi control strain with increasing concentrations of the various antibiotics, see **Fig. S3**. (**E**) The AlphaFold 3-predicted structure of CxaR, with its three proposed anti-parallel sets of TPR-like helices colored red, blue, and purple. These helices form a slightly concave surface groove, as indicated in the figure

Strikingly, and in line with the results from the CRISPRi-seq screen, subinhibitory concentrations of penicillin G rescued the growth defects of the *cxaR* knockdown and deletion mutants. When treated with penicillin G, the mutants grew similarly to the controls, both in liquid and on solid medium (**Fig 1**, **S3A Fig**, **S5A Fig**). The minimal inhibitory concentration (MIC) of the *cxaR* mutants, however, remained unaltered (**S3A Fig**).

To determine whether the penicillin G-mediated growth stimulation was a general response to growth inhibition or specific to certain antibiotics, the JE2 *cxaR* knockdown strain was tested against a panel of antibiotics, targeting different cellular processes (**Fig 1D** and **S3 Fig**). Subinhibitory concentrations of all tested β-lactams (penicillin G, oxacillin, and cefoxitin) fully rescued growth of CxaR depleted cells (**Fig 1D**, **S3A-C Fig**). Similarly, other antibiotics targeting the cell wall, such as moenomycin (inhibiting transglycosylation activity) and tunicamycin (inhibiting early stages in wall teichoic acid and peptidoglycan biosynthesis), also fully restored growth (**Fig 1D**, **S3G-H Fig**). Partial growth rescue was observed for bacitracin (inhibiting lipid carrier recycling) and teicoplanin (glycopeptide inhibiting cell wall synthesis by binding to the D-Ala-D-Ala terminus of peptidoglycan precursors), while another glycopeptide with the same mechanism of action, vancomycin, did not seem to stimulate growth (**Fig 1D**, **Fig S3D-F**). Furthermore, translation inhibitors binding to the 30S ribosomal subunit, including kanamycin and streptomycin, as well as the DNA gyrase inhibitor novobiocin, partially restored growth of the CxaR depleted strain (**Fig 1D**, **S3N,O,J Fig**). On the other hand, the tested antibiotics targeting topoisomerase IV (ciprofloxacin), folate biosynthesis (trimethoprim), transcription (rifampicin), and the 50S ribosomal subunit (erythromycin) did not stimulate growth at any concentration (**Fig 1D**, **S3I,K,L,M Fig**).

### CxaR is conserved among *Staphylococcaceae* and has structural resemblance to TPR motifs

*cxaR* is part of the *S. aureus* core genome, and phylogenetic analysis of CxaR homologues from 55 different species within the *Staphylococcaceae* family demonstrated a high degree of conservation among the homologs from the closely related genera *Staphylococcus*, *Macrococcus*, and *Mammaliicoccus*, with 44 out of 165 amino acids being completely invariable (**S4A-B Fig**). *cxaR* does not encode any signal peptide nor any transmembrane helices. The high-confidence AlphaFold-predicted structure of CxaR consists of two short N-terminal α-helices, six similarly sized antiparallel α-helices forming a slightly concave groove, and finally a longer C-terminal α-helix protruding from the rest of the structure (**Fig 1E**). When mapping residue conservation onto the predicted structure of CxaR, we found that highly conserved residues were primarily located at positions mediating hydrophobic interactions between the antiparallel α-helices and within the groove (**Fig 1E**, **S4 Fig**). HHpred [61] and HMMER [62] were used to identify potential protein domains of CxaR. Interestingly, both tools revealed a substantial number of significant hits with tetratricopeptide repeat (TPR) motifs. TPR motifs consist of 34-residue antiparallel helix pairs, typically repeated 3 to 16 times to form grooved, amphipathic binding surfaces that facilitate protein-protein interactions [63]. Although the sequences of TPR motifs are highly variable and therefore at times difficult to detect, they are involved in a wide range of cellular processes [64, 65]. The most probable TPR helix pairs of CxaR, along with the resulting binding groove, are labelled in **Fig 1E**.

### CxaR affects splitting of daughter cells

To better understand the growth phenotypes of the *cxaR* mutants, phase contrast and epifluorescence microscopy were performed on samples taken at different stages of growth (indicated in **Fig 2A**). Cells lacking CxaR clearly displayed increased lytic activity. Even before the drop in OD, a noticeably higher proportion of mutant cells appeared lysed compared to wild-type cells (hour 6 and 8, **Fig 2B and D**, **S5 Fig**). While the morphology of the live mutant cells did not significantly diverge from the wild-type, including comparable cell size (**Fig 2C**), we often observed *cxaR* mutant cells with a nick at septum displaying DNA dispersal, as visualized by DAPI-staining (**Fig 2B and D**). After the temporal decline in OD, few intact cells remained, and a large amount of cellular debris was observed (**Fig 2B**). During the subsequent growth recovery phase, less lysis was observed. Strikingly, addition of penicillin G to cells lacking CxaR significantly reduced lysis at all recorded time points, with a particularly notable reduction at the 10-hour mark, when the majority of CxaR-deficient cells would otherwise have lysed (**S5 Fig**).

**Fig. 2.**
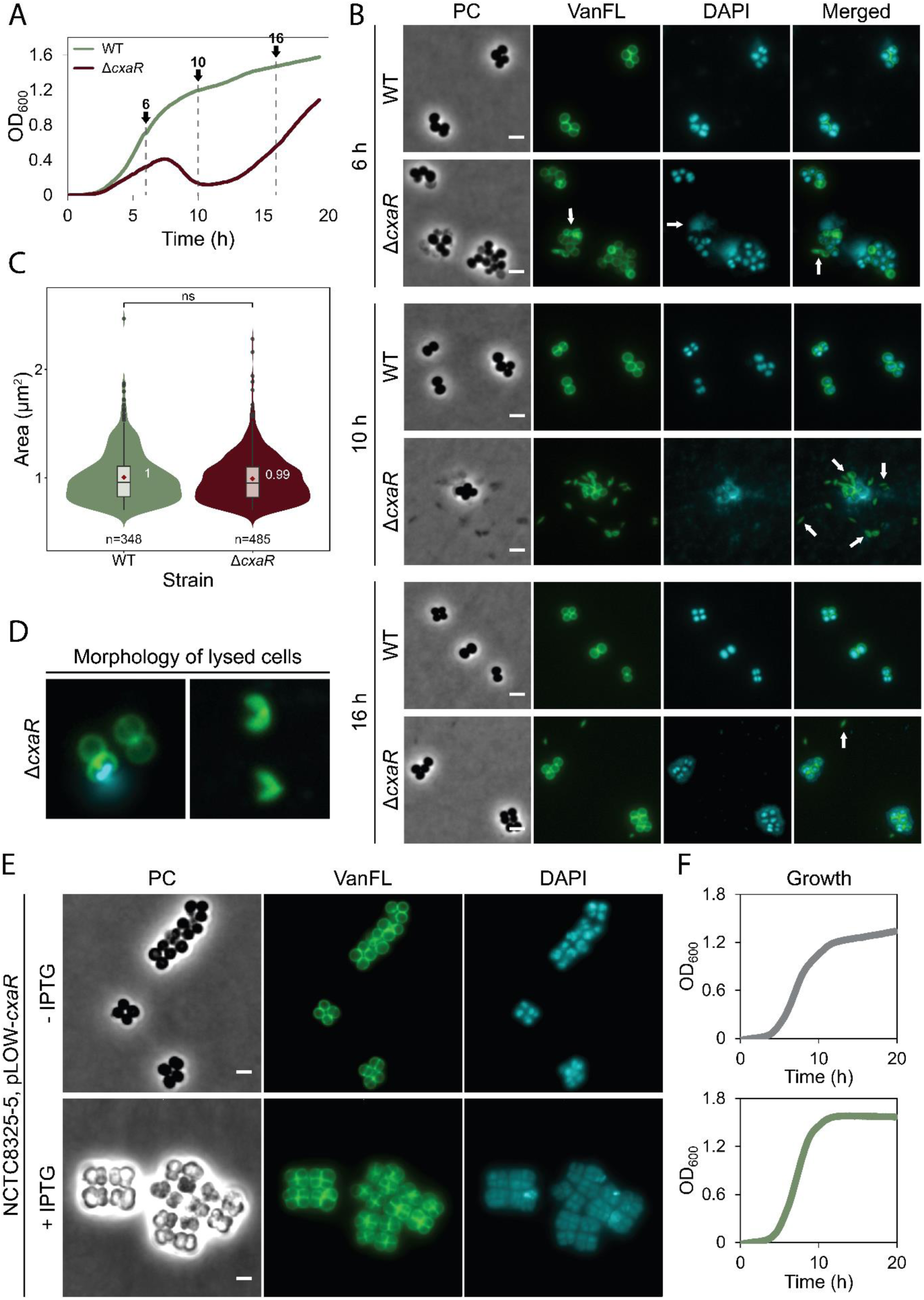
*cxaR* deletion increases lytic activity, while overexpression impairs cell division. (**A**) Growth curves of NCTC8325-4 Δ*cxaR* and wild-type, with the timepoints for cell sampling used in microscopy (shown in panel **B**). The graphs represent averages from triplicate measurements. (**B**) Representative phase-contrast and fluorescence micrographs of Δ*cxaR* and wild-type cells stained with VanFL (cell wall) and DAPI (nucleoid) after 6, 10, and 16 hours of incubation at 37°C. White arrows point to cells with peptidoglycan hydrolysis (VanFL), nucleoid dispersal (DAPI), and anucleate cells (merged). The scale bars are 2 µm. (**C**) Violin plots showing the cell area distributions (in µm^2^) for wild-type (1.00 ± 0.24 µm^2^) and Δ*cxaR* (0.99 ± 0.22 µm^2^) after 6 hours at 37°C, determined using MicrobeJ. No significant difference in size was detected between the strains (ns, *P*-value > 0.05, Mann-Whitney U test). The number of cells analyzed is indicated in the figure. (**D**) Magnified merged fluorescence images showing the predominant lytic phenotypes of the Δ*cxaR* cells. Cells commonly exhibited a nick at septum accompanied by DNA dispersal at the earlier timepoints, progressing to crescent-shaped debris by the 10-hour mark, indicative of widespread lysis. (**E**) Representative phase-contrast and fluorescence micrographs of NCTC8325-4 cells carrying the pLOW-*cxaR* plasmid for inducible overexpression of CxaR, stained with VanFL (cell wall) and DAPI (nucleoid). Cells were grown at 37°C with or without 500 µM IPTG until an OD_600_ of approximately 0.4 was reached. Upon induction, cells displayed clear separation defects, including cell clumping and enlarged cells with multiple septa. The scale bars are 2 µm. (**F**) Growth curves of NCTC8325-4 cells with the pLOW-*cxaR* plasmid, grown in the absence (gray) or presence (green) of 500 µM IPTG for CxaR overexpression. The graphs represent averages from triplicate measurements.

CxaR overexpression was also analyzed (**Fig 2E-F**). Elevated levels of CxaR did not result in any growth deficiency. In fact, these cells reach a slightly higher final OD compared to those with normal CxaR levels (**Fig 2F**). Morphologically, cells with excess CxaR clustered together, frequently forming tetrads, where a second round of division is initiated before the previous pair of daughter cells have separated (**Fig 2E**). Notably, very few of the CxaR overexpressing cells were observed without a septum, indicating that excess CxaR does not impair septum synthesis but rather disrupts the separation of daughter cells after septum completion.

The lysis phenotype was further investigated by transmission electron microscopy (TEM). As expected, TEM analysis of CxaR depleted cells also revealed a significantly higher proportion of lysed cells compared the control strain (**Fig 3A and C**). Consistent with earlier observations, penicillin-treated *cxaR* samples showed a drastic reduction in lysis compared to the untreated samples (**Fig 3A and C**). Interestingly, the mutant cells frequently displayed premature splitting of daughter cells, where the septum appeared to be degraded before it was completed (marked with red asterisks in **Fig 3A and G-I**). Variations of this phenotype were observed, with some cells exhibiting a nick prior to or at early stages of septal growth (**Fig 3G**), while others underwent splitting just before septum completion (**Fig 3H**). Some cells also underwent splitting at both ends of the septum simultaneously (**Fig 3I**). Furthermore, while most live *cxaR* cells resembled the control cells (**Fig 3D-E**), a subset exhibited a significantly thicker cell wall (**Fig 3F**), potentially a strategy adapted by some mutant cells to evade lysis. Taken together, these findings indicate that the lysis observed in cells lacking CxaR is due to premature septum splitting, and that penicillin G mitigates this by preventing spontaneous lysis in the mutant.

**Fig. 3.**
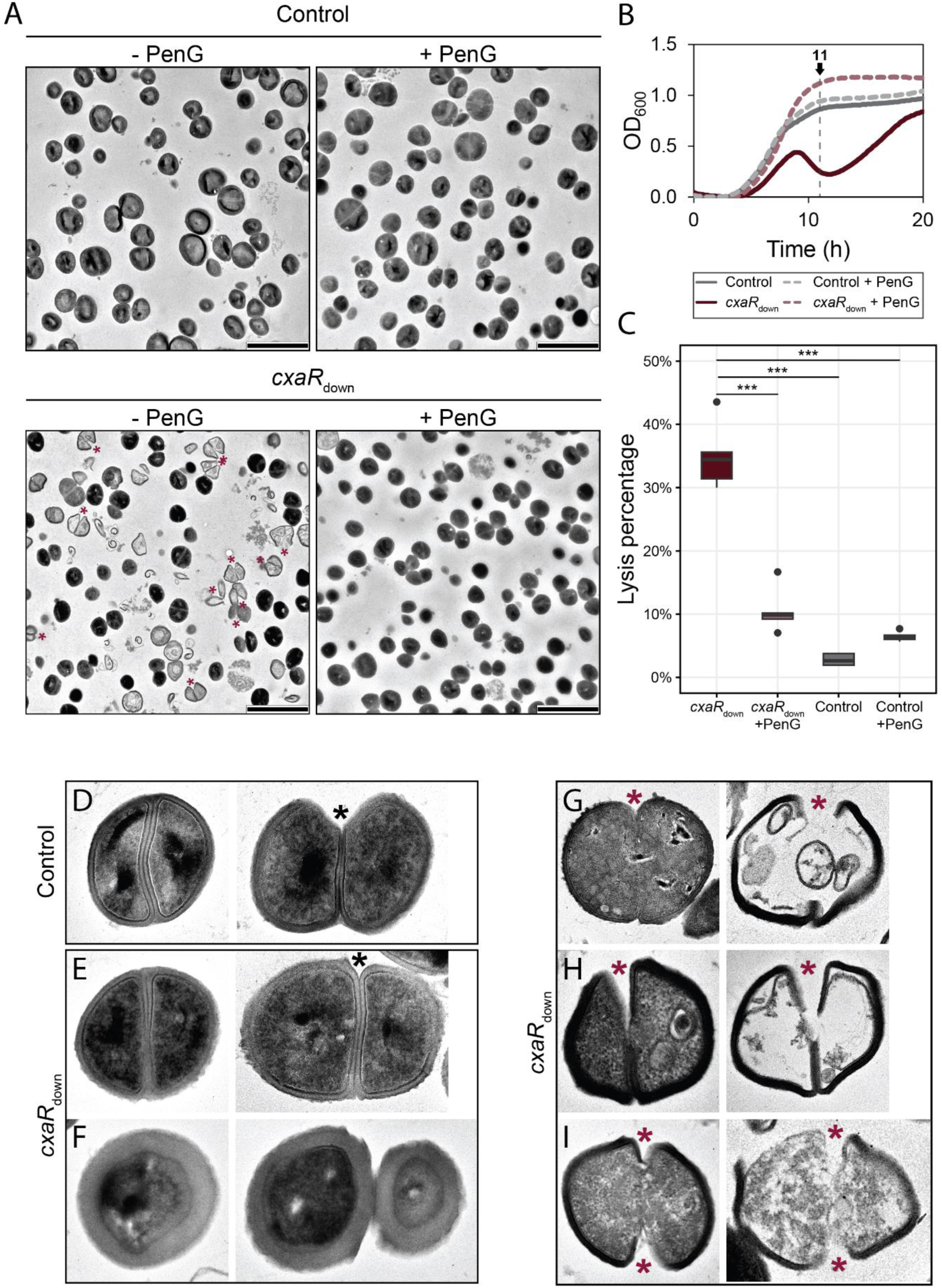
The increased lysis observed in the absence of CxaR is due to premature splitting of daughter cells, a phenomenon significantly mitigated by penicillin G treatment. (**A**) TEM images of the NCTC8325-4 *cxaR* knockdown mutant (*cxaR*_down_) and CRISPRi control strain grown in TSB for 11 hours at 37°C, in the absence or presence of sublethal concentrations of penicillin G (1 ng/mL). Red asterisks mark cells that have undergone cell splitting prior to septum completion. The scale bars are 2 µm. (**B**) Growth curves of NCTC8325-4 *cxaR*_down_ and CRISPRi control, with (dashed line) or without (solid line) sublethal concentrations of penicillin G (1 ng/mL), showing the sampling timepoint for TEM imaging. The graphs represent averages from triplicate measurements. (**C**) Box plots showing the proportion of lysed cells, as observed by TEM, that retained minimal structural integrity. Significant differences between the strains are indicated with asterisks (***, *P*-value < 0.0005, Tukey’s Honest Significant Difference test). The number of cells analyzed, from left to right, is 231, 303, 241, and 250. (**D-I**) Representative TEM images of CRISPRi control and *cxaR*_down_ cells, highlighting their characteristic morphologies. (**D**) Control cells displaying normal coccoid morphology. *cxaR* knockdown cells displaying: (**E**) wild-type-like morphology, (**F**) significantly thicker cell wall, (**G**) a septal nick prior to or at early stages of septal growth, (**H**) premature daughter cell splitting occurring just before septum completion, and (**I**) simultaneous splitting at both ends of the forming septum. Black asterisks mark cells in which hydrolysis of the peripheral septal wall occurred after septum completion, while red asterisks mark cells initiating daughter cell separation, despite an incomplete septum.

### Reduced Sle1 activity can compensate for the lack of CxaR

Since cell splitting in *S. aureus* is known to be temperature-responsive [20], we examined the growth of *cxaR* mutant cells across a range of temperatures. The mutants exhibited a clear temperature-dependent growth phenotype (**Fig 4A**, **S6 Fig**); the growth defect was exaggerated at 30°C, while it was recovered at higher temperatures (42-45°C). Notably, when plated at 30°C, the *cxaR* deletion mutant displayed heterogeneous colony size, with both small and large-sized colonies, in contrast to the uniform, large colony size of the wild-type (**Fig 4B**). The colony size variation also appeared at 37°C, although to a lesser extent (**Fig 4A**, **S6B Fig**). We hypothesized that the larger sized colonies at 30°C had obtained suppressor mutations. Five Δ*cxaR* colonies were picked and shown to exhibit wild-type like growth patterns (**Fig 4B and 4C**). Whole-genome re-sequencing of these mutants revealed that each had independently acquired additional mutations associated with the PGH *sle1* (**S2 Table**). Two of the suppressor mutants had substitutions in the catalytically active CHAP-domain, responsible for the hydrolytic activity of Sle1 (M292I and G315C), two resulted in truncated versions of Sle1 (Q197* and V5*), while the last one acquired an alteration in the Shine-Dalgarno sequence of *sle1* (G>A in the RBS). The mutations thus likely resulted in inactivation and/or reduced activity of Sle1 (see below).

**Fig. 4.**
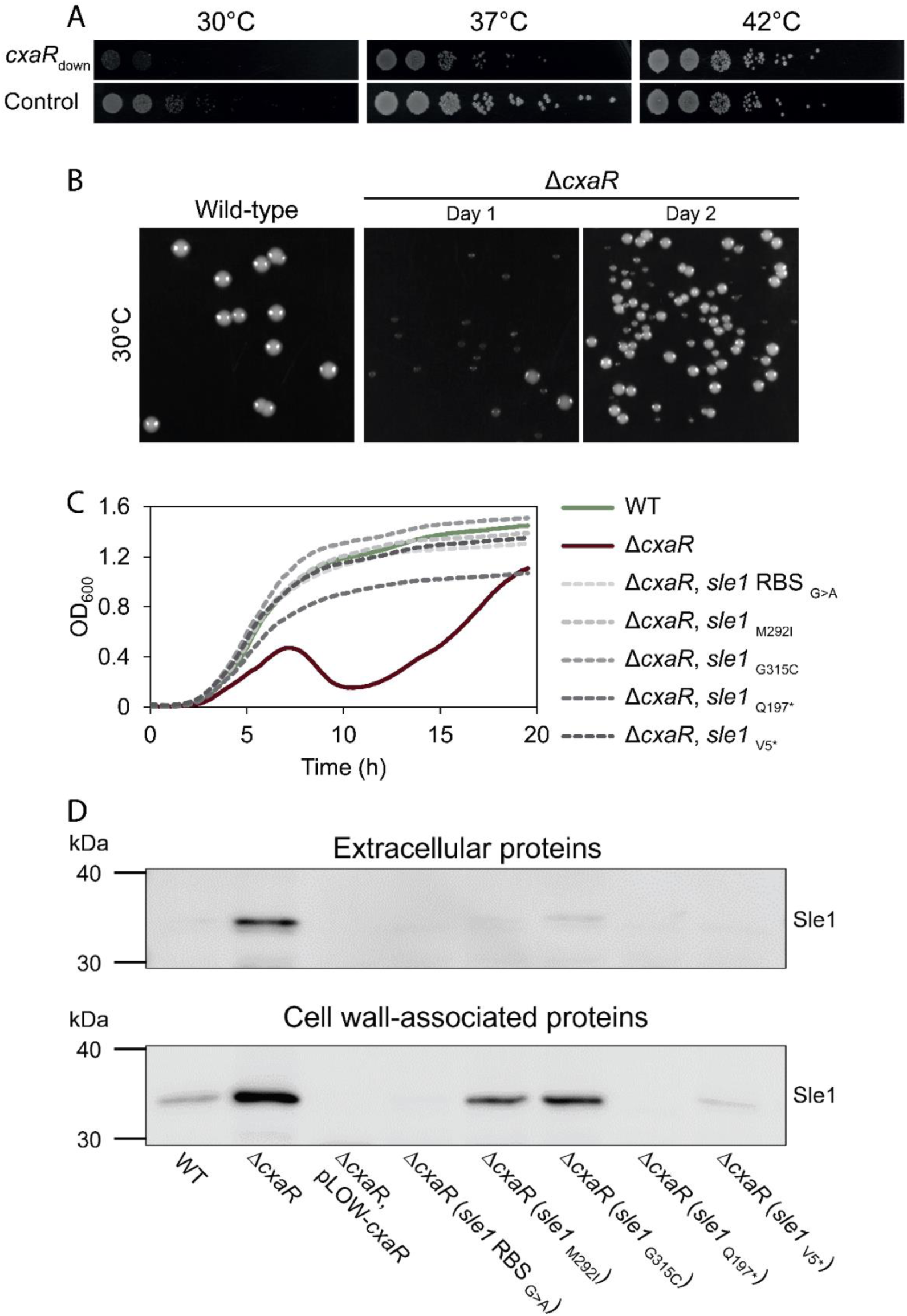
Spontaneous *sle1*-associated mutations frequently rescued the defects of the *cxaR* deletion mutant at 30°C. (**A**) Spotting assays comparing growth of the NCTC8325-4 *cxaR* knockdown mutant (*cxaR*_down_) and CRISPRi control at 30°C, 37°C, and 42°C. The bacterial cultures were grown at 37°C to an OD_600_ of 0.5, 10-fold serially diluted, and 5 µL of each dilution was spotted onto TSA plates that were incubated at the indicated temperatures. Compared to the control, the *cxaR*_down_ mutant was cold-sensitive and heat-stable. (**B**) At 30°C, the NCTC8325-4 Δ*cxaR* mutant formed both very small colonies and larger suppressor colonies, unlike the homogeneous colony size observed for wild-type. When overnight cultures of Δ*cxaR* grown at 37°C were plated on TSA and incubated at 30°C, some larger colonies appeared among the very small ones (day 1). When single colonies from these plates were grown overnight at 30°C and replated on TSA at 30°C, the plates exhibited predominantly large suppressor colonies (day 2). (**C**) Growth of five independently isolated large Δ*cxaR* colonies, all found to have different *sle1*-associated mutations, in TSB at 37°C. All five suppressor mutants displayed growth patterns more closely resembling the wild-type than the original Δ*cxaR* strain. The graphs represent averages from triplicate measurements. (**D**) Western blot analysis of Sle1 levels in the extracellular and cell wall-associated protein fractions of the Δ*cxaR* suppressors with *sle1*-associated mutations. The Sle1 levels of the suppressor mutants were compared to those of wild-type, *cxaR* deletion, and *cxaR* overexpressing cells.

To further confirm that the growth defects caused by a lack of CxaR can be rescued by the absence of *sle1*, *sle1* was knocked down in the Δ*cxaR* strain (**S7 Fig**). As expected, depletion of Sle1 in the *cxaR* mutant resulted in loss of the biphasic growth and thus a reversion to a wild type-like growth phenotype (**S7 Fig**).

### Sle1 levels are highly elevated in the absence of CxaR and reduced upon CxaR overexpression

To investigate the apparent functional link between CxaR and Sle1 levels, we performed Western blot analyses using an antibody that recognizes Sle1 (35.8 kDa) [20]. Strikingly, a clear accumulation of Sle1 was detected in cells lacking CxaR (Δ*cxaR* and *cxaR*_down_ in **Fig. 5, S8 Fig, S9 Fig**). Sle1 accumulation was observed both in the cell wall fraction, the extracellular fraction, and the whole cell lysate. In contrast, when CxaR is overexpressed, Sle1 completely disappears in all fractions (**Fig. 5, S8 Fig, S9 Fig**). Thus, the cellular levels of Sle1 are controlled by CxaR, as both its presence and absence directly correlate with Sle1 protein levels.

**Fig. 5.**
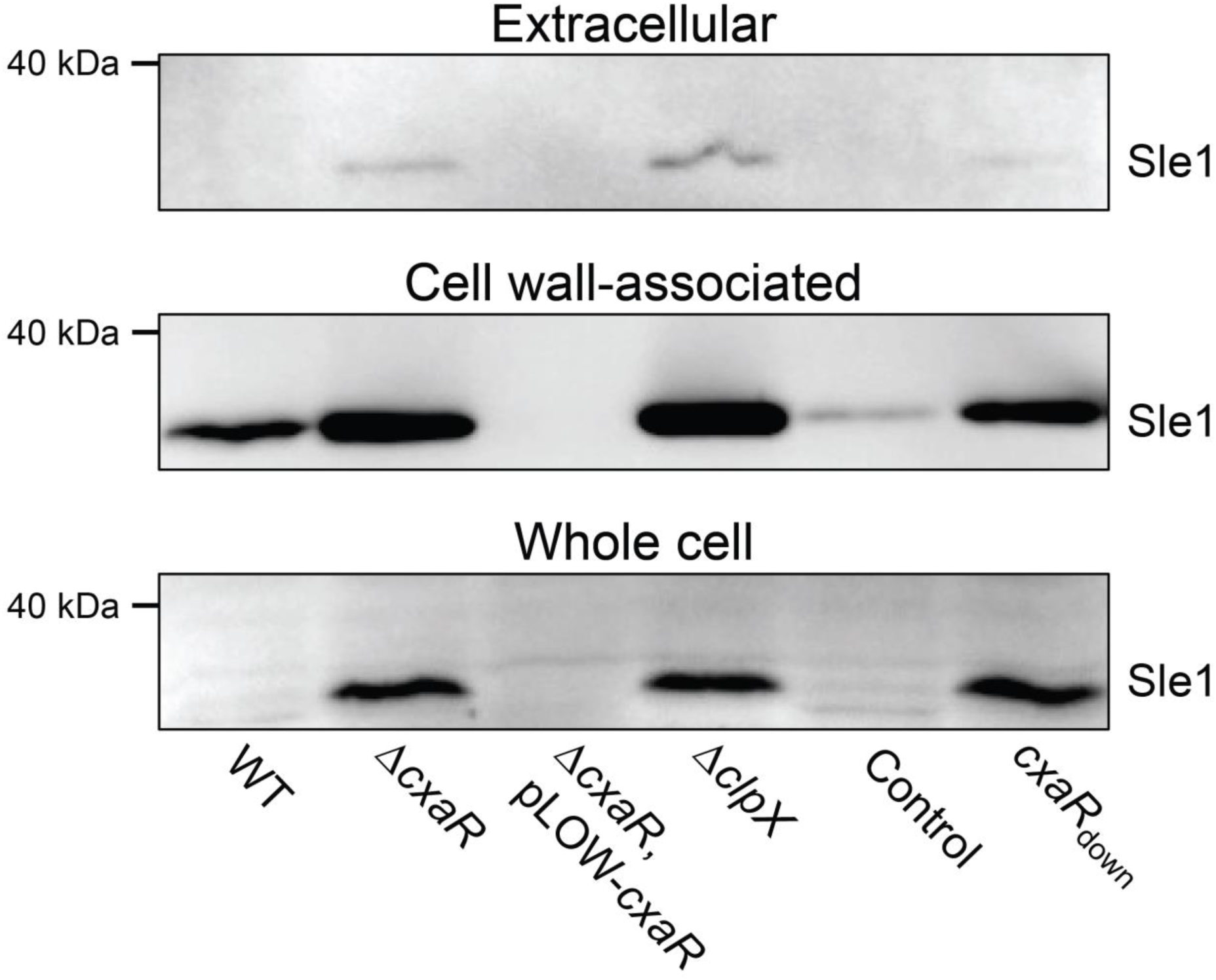
Sle1 accumulates in the absence of CxaR, while overexpression of CxaR causes Sle1 depletion below detectable levels. Western blot analysis of Sle1 levels in cell wall-associated protein fractions, extracellular protein fractions, and whole-cell lysates using an anti-Sle1 antibody. The blots show that Sle1 levels were elevated in NCTC8325-4 cells lacking CxaR (Δ*cxaR* and *cxaR*_down_) or ClpX (Δ*clpX*), while Sle1 was depleted in NCTC8325-4 cells overexpressing CxaR (Δ*cxaR*, pLOW-*cxaR*), compared to the wild-type and CRISPRi control cells.

We then performed immunoblot assays with cell wall-associated and extracellular protein extracts from the *cxaR* suppressor mutants (**Fig 4D**). This revealed reduced levels of cell wall-associated Sle1 in cells with the *sle1* RBS mutation as well as in those encoding truncated Sle1 versions, compared to the wild-type. The mutants with single CHAP-domain substitutions also exhibited slightly reduced Sle1 levels, although not restored to wild-type levels. This further supports the notion that reducing Sle1 levels or activity can compensate for the loss of CxaR.

Given that reduced levels of Sle1 seem to rescue growth of cells lacking CxaR, we investigated whether the same mechanism was at play when growth was restored at high temperatures and/or with the addition of penicillin G. Indeed, both treatment with penicillin G and incubation at 42°C reduced Sle1 levels, both in wild-type cells and cells without CxaR (**S10A-B Fig**). We also tested whether the other antibiotics that stimulated growth of the *cxaR*_down_ mutant had a similar effect on Sle1 levels (**Fig 1D**). Indeed, a reduction in cell wall-associated Sle1 was observed, although the extent of reduction varied. Oxacillin, bacitracin, moenomycin, tunicamycin, and kanamycin all appeared to decrease Sle1 levels in the Δ*cxaR* strain to some degree. However, for teicoplanin and novobiocin, the effect on Sle1 levels was less evident (**S10C Fig**), consistent with their comparable limited ability to rescue growth of the *cxaR*_down_ mutant. Together, this suggests that the stimulatory effects of antibiotics observed for the Δ*cxaR* strain at least partially can be explained by reduced levels of Sle1.

### CxaR regulates Sle1 levels in a ClpXP-dependent manner

Previous studies have shown that the cellular levels of Sle1 are regulated by the ClpXP protease complex [38, 39], and similarly, we observed a strong accumulation of Sle1 Δ*clpX* cells and cells devoid of ClpXP complex formation (ClpX_I265E_) (**Fig 5**, **Fig 6A**, **S9 Fig**). In fact, *cxaR* mutant cells phenocopies *clpX* mutants to a large degree, including the observed temperature-dependent growth defect and growth rescue by cell wall targeting antibiotics (**S11 Fig**) [39, 41, 44]. These parallels suggest that CxaR plays a role in the ClpXP-mediated Sle1 degradation. Notably, the level of Sle1 accumulation in cells lacking either ClpXP activity or CxaR was comparable (**Fig 5**, **S9 Fig**). If CxaR was primarily involved in regulating *sle1* expression, one would expect the combined loss of CxaR and ClpXP to produce a significantly stronger Sle1 accumulation compared to either single mutant. However, ClpX_I265E_ cells with CxaR depletion showed no greater Sle1 accumulation than ClpX_I265E_ cells (**S9 Fig**). In addition, there was no reduction in cell wall-associated Sle1 when CxaR was overexpressed in the Δ*clpX* mutant, indicating that the mechanism in which CxaR controls Sle1 levels is not active when *clpX* is deleted (**S9 Fig**). Furthermore, a *cxaR clpX* double knockdown strain did not exacerbate the phenotypes of the two single knockdowns (**S11A and C Fig**). Together, this suggests that CxaR and ClpXP likely operate within a shared pathway rather than through independent mechanisms.

**Fig. 6.**
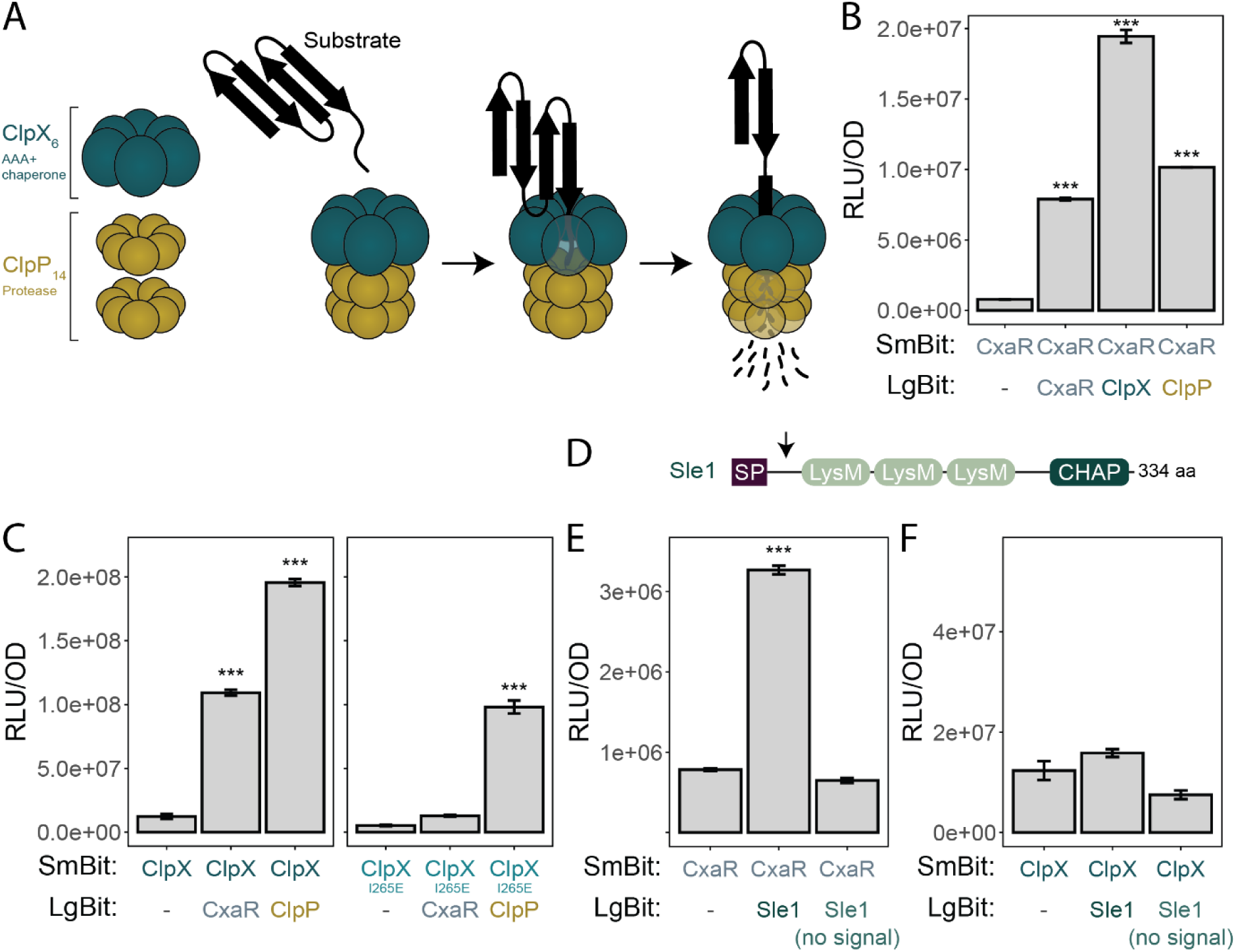
CxaR interacts with ClpX and Sle1. (**A**) Schematic overview of a single-capped ClpXP structure and its function. Single-capped ClpXP consist of a hexameric ClpX ATPase ring and two heptameric rings of the ClpP peptidase, forming a proteolytic complex. ClpX recognizes and unfolds protein substrates, either directly or via adaptor proteins, and translocates them through its central pore into the proteolytic chamber of ClpP, where they are degraded into small peptide fragments, ensuring selective and irreversible degradation of proteins in the cell. (**B-C**) Split luciferase assays identifying potential protein-protein interactions between (**B**) CxaR and CxaR/ClpX/ClpP, and (**C**) ClpX/ClpX_I265E_ and CxaR/ClpP. Bars represent mean luminescence (RLU/OD_600_) from four replicates, with error bars indicating the standard deviation. Significant interactions were assessed with one-way ANOVA followed by Tukey’s Honest Significant Difference test (***, *P*-value < 0.001). CxaR displayed self-interaction and interactions with both ClpX and ClpP. Similarly, ClpX interacted with CxaR and ClpP, however the interaction with CxaR appears to be lost when ClpX does not form a complex with ClpP. (**D**) The modular structure of the Sle1 amidase, consisting of a N-terminal signal peptide (SP) for secretion, three cell wall-binding LysM domains, and an enzymatically active histidine-dependent amidohydrolase/peptidase (CHAP) C-terminal domain. The signal peptide of Sle1 is post-translationally processed, as indicated by the arrow. (**E-F**) Split luciferase assays assessing potential protein-protein interactions between (**D**) CxaR and Sle1 (with and without the signal peptide), and (**E**) ClpX and Sle1 (with and without the signal peptide). Bars represent mean luminescence (RLU/OD_600_) from four replicates, with error bars indicating the standard deviation. Significant interactions were assessed with one-way ANOVA followed by Tukey’s Honest Significant Difference test (***, *P*-value < 0.001). CxaR showed interaction with the precursor of Sle1 (with signal peptide), while ClpX did not interact with the precursor nor the mature form of Sle1.

### CxaR directly interacts with ClpXP and Sle1

Seeing that structural modeling of CxaR suggests it may contain a TPR-like domain (**Fig 1E**) commonly associated with protein-protein interactions [64, 66, 67], we investigated the potential role of CxaR in mediating interactions between Sle1 and ClpXP. We employed the split luciferase system [68], where the proteins of interest are conjugated to the complementary small (SmBit) and large (LgBit) subunits of luciferase, which produce a quantifiable luminescence signal if they come into close proximity when expressed in *S. aureus*.

Initially, CxaR self-interaction was tested, as TPR-containing proteins commonly exhibit self-oligomerization when involved in regulatory roles [69, 70]. Indeed, CxaR appears to oligomerize, as that the self-interaction displayed a significant 10-fold higher value than the control (**Fig 6B**). Next, interactions between CxaR, ClpX, and ClpP were tested. Luminescence readings between all three were highly significant (**Fig 6B-C**). While it is well established that ClpX and ClpP form a proteolytic complex (**Fig 6A and C**), CxaR also appears to interact with both proteins, displaying a 25-fold and 13-fold increase in signal compared to the control, respectively (**Fig 6B**). However, the interaction between CxaR and ClpX appears to be lost when ClpX contains the I265E mutation (**Fig 6C**), suggesting that CxaR only interacts with the ClpXP complex. Notably, the interaction of CxaR with ClpX generated a stronger signal than the interaction with ClpP.

Moreover, CxaR also displayed a significant interaction with the precursor of Sle1, with a 4-fold increase in luminescence signal compared to the control (**Fig 6D-E**). Interestingly, however, this interaction was not observed when Sle1 was expressed without the signal peptide (**Fig 6D-E**). Indeed, in the AlphaFold3 predicted interaction model, supported by a high interaction score, the first LysM domain of Sle1 (located in the N-terminal part close to the signal sequence) interacts with the predicted binding groove of CxaR (**Fig 1E**, **S12 Fig**). Neither the precursor nor the mature form of Sle1 showed any detectable interaction with ClpX (**Fig 6F**). Together this strongly suggests that CxaR functions as an adaptor or intermediary to mediate the interaction between Sle1 and the ClpXP complex, facilitating Sle1 degradation. Since the Sle1 precursor is transported across the cytoplasmic membrane via its N-terminal signal peptide (**Fig 6D**), it is expected that only the precursor of Sle1 is accessible for degradation by the cytoplasmic ClpXP complex.

### CxaR localization is ClpX-dependent

ClpX has been shown to localize in single foci near the membrane, positioned close to the ingrowing septum, in *S. aureus* cells undergoing septation [45]. This was confirmed here by expressing an inducible ClpX-mCherry fusion (**S13A Fig**). Furthermore, a CxaR-GFP fusion appeared to localize at midcell in dividing cells (**Fig 7A**, **S13B-C Fig**). Indeed, when co-expressing ClpX-mCherry and CxaR-GFP, a clear co-localization of the two proteins was observed, with their fluorescent signals appearing at the exact same positions in approximately 92% of cells (n = 635) (**Fig 7C and D**). To determine whether CxaR depends on ClpX for its localization, a plasmid encoding CxaR-GFP was transformed into the Δ*clpX* mutant. In these cells, the distinct signal pattern was lost (**Fig 7F**), suggesting that CxaR requires ClpX for its spatiotemporal localization. However, when ClpX-mCherry was introduced into Δ*cxaR* cells, the localization remained unchanged (**Fig 7E**), indicating that ClpX localization is independent of CxaR. It should be noted that while the CxaR-GFP fusion seems to be functional in terms of growth (**Fig 7B**), cells expressing CxaR-GFP, either from its native promoter on the chromosome or from a plasmid, exhibited cell separation defects, including cell clumping (**Fig 7A**, **S13B-C Fig**).

**Fig. 7.**
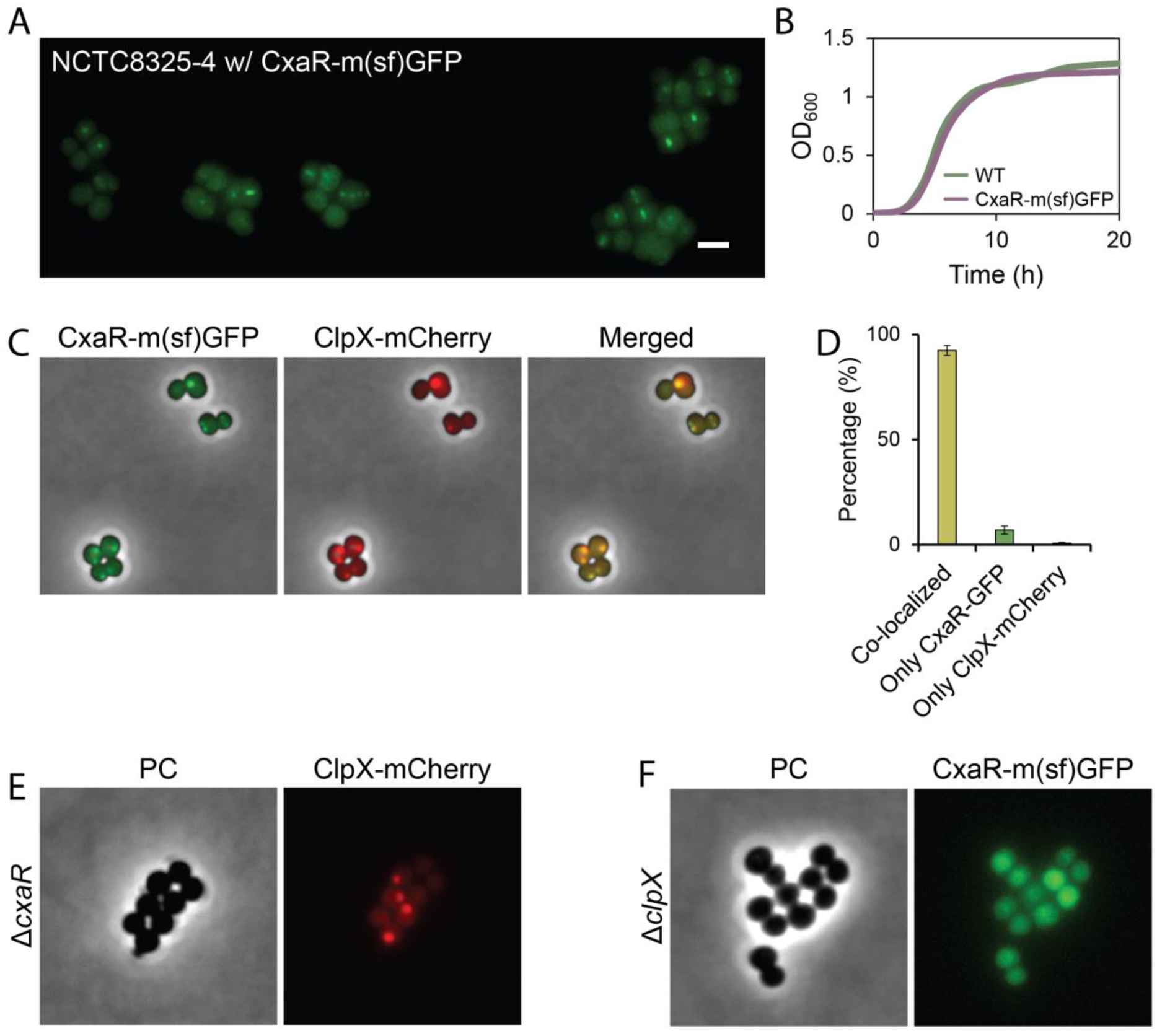
CxaR localizes adjacent to the septum in a ClpX-dependent manner. (**A**) Fluorescence microscopy of the NCTC8325-4 strain expressing chromosomally integrated CxaR-GFP under its native promoter, showing that CxaR-GFP localizes at midcell in dividing cells. The scale bar represents 2 µm. (**B**) Growth curves of NCTC8325-4 wild-type and chromosomally expressing CxaR-GFP cells in TSB at 37°C, demonstrating that GFP tagging of CxaR does not impair growth. The graphs represent averages from triplicate measurements. (**C**) Fluorescent microscopy of NCTC8325-4 cells co-expressing CxaR-GFP (from the chromosome) and ClpX-mCherry (from a plasmid induced with 80 μM IPTG), showing clear co-localization of the two proteins. (**D**) Quantification of CxaR-GFP and ClpX-mCherry co-localization in NCTC8325-4 cells expressing both fusion proteins. Co-localization was analyzed by manual examination of the fluorescence signals in 300– 350 randomly selected cells from two biological replicates (n = 635). CxaR-GFP and ClpX-mCherry were co-localized in 92.4 ± 2.4% of cells, 6.9 ± 1.9% of cells displayed only CxaR-GFP signal, and 0.6 ± 0.5% showed only ClpX-mCherry, confirming spatiotemporal co-localization of the two proteins during cell division. (**E-F**) Fluorescence microscopy of (**H**) Δ*clpX* cells expressing CxaR-GFP from a plasmid and (**I**) Δ*cxaR* cells expressing ClpX-mCherry from a plasmid. Expression of the plasmids were induced with the addition of 10 μM and 160 uM IPTG, respectively. In the absence of ClpX, CxaR-GFP loses septal-adjacent localization, indicating that CxaR requires ClpX for its spatiotemporal localization. In contrast, ClpX-mCherry retained its focal localization pattern in Δ*cxaR* cells, demonstrating that ClpX localization does not depend on the presence of CxaR.

## Discussion

In this work, we have discovered a novel factor critical for regulating cell splitting in *S. aureus*. Our results demonstrate that the hitherto unstudied protein CxaR acts as an adaptor protein, controlling the levels of Sle1, a PGH crucial for staphylococcal cell separation. Specifically, our model proposes that CxaR binds the Sle1 precursor prior to its export and translocates it to the ClpXP protease for degradation (**Fig 8**).

**Fig. 8.**
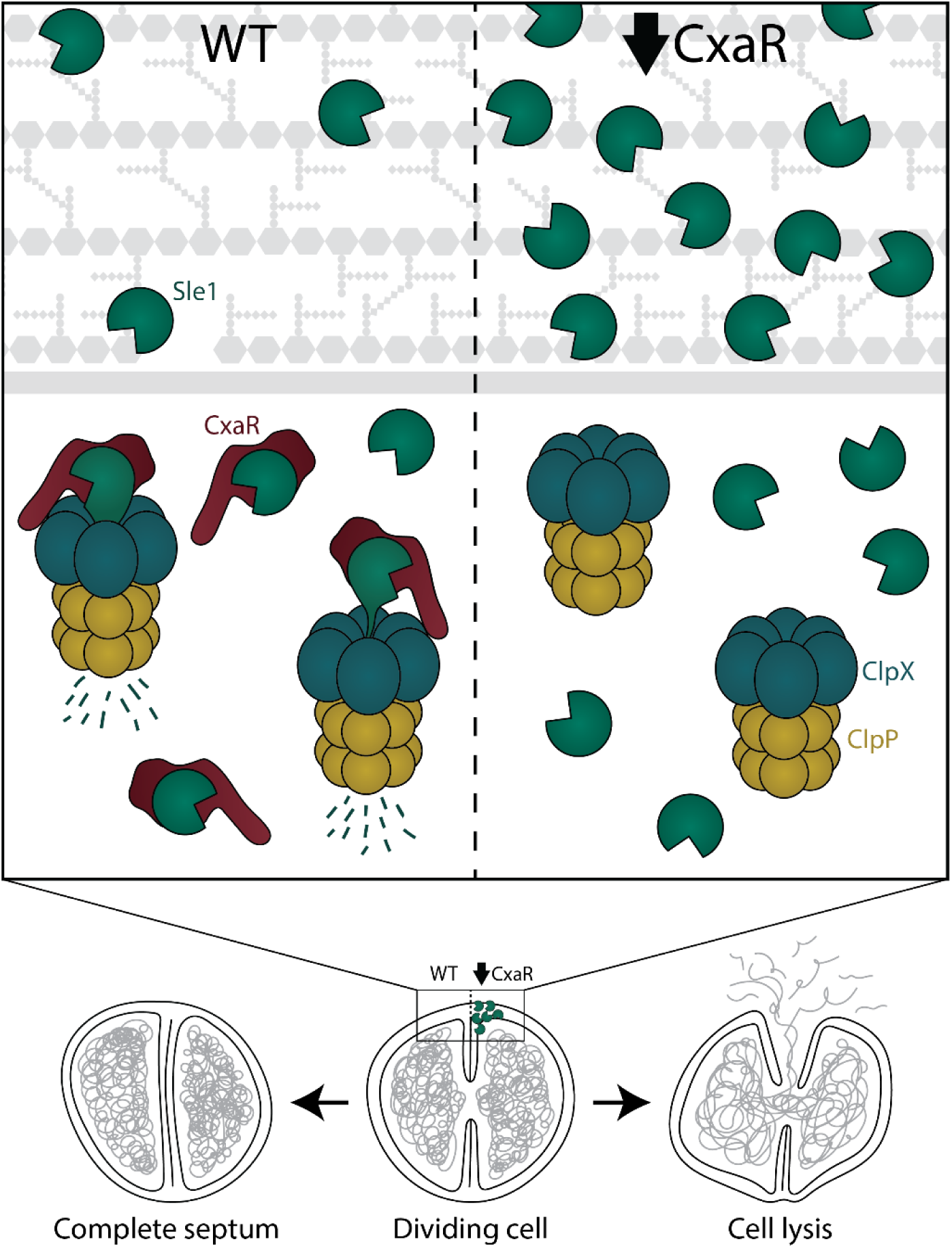
CxaR facilitates ClpXP-mediated degradation of Sle1, ensuring proper separation of staphylococcal daughter cells. Under normal conditions, CxaR regulates Sle1 levels by delivering its precursor to ClpXP for degradation, ensuring that cell splitting is triggered only after septum completion. In the absence of CxaR, ClpX and ClpP still form a proteolytic complex capable of degrading various proteins, but not Sle1, as its mode of delivery is missing. Consequently, Sle1 accumulates to high levels, causing hydrolysis of the incomplete septum and subsequent cell lysis.

This model is supported by several experimental observations. First, we observed that *cxaR* mutants phenocopies *clpX* mutants in a number of aspects, including biphasic growth profiles, temperature-dependent growth phenotypes with exacerbation of the growth defect at suboptimal and improved growth at supraoptimal temperatures [44], high lysis frequency due to premature cell splitting [41], and accumulation of Sle1 [38]. These defects were found to be alleviated by certain antibiotics, including β-lactams and tunicamycin, as well as inactivation of Sle1, similar to what has previously been observed for cells lacking ClpX [41]. Consistently, overexpression of CxaR led to depletion of cellular Sle1 due to enhanced degradation, causing severely impaired cell splitting. Second, split luciferase assays, used to assess protein-protein interactions *in vivo*, showed that CxaR interacts directly with the ClpXP complex, as well as Sle1. However, no direct interaction between Sle1 and ClpX was detected in these assays, suggesting that an intermediate protein facilitates the delivery of Sle1 to ClpXP for targeted degradation. Third, subcellular localization analysis showed that CxaR co-localizes with ClpX near the septum in dividing *S. aureus* cells. Additionally, the localization of CxaR was dependent on the presence of ClpX, indicating that its function is closely linked to ClpX and likely cell division. Moreover, it should also be noted that CxaR exhibited the highest score among identified staphylococcal ClpXP substrates when a proteolytically inactive ClpP variant (ClpP^trap^) was expressed *in vivo* in *S. aureus* [38]. However, CxaR-GFP does not appear to accumulate in *S. aureus* cells lacking ClpXP activity (**S13D Fig**), suggesting that its retention in the ClpP^trap^ is likely due to other interactions than direct degradation. Finally, structural analyses revealed that CxaR resembles proteins containing TPR motifs, which are typically involved in facilitating protein-protein interactions [64, 66, 67]. This structural similarity suggests that CxaR may bind specific substrates to mediate their interactions or delivery. Thus, dysregulation of CxaR levels have detrimental effects on cell splitting and growth; Sle1 accumulates in the absence of CxaR, resulting in cell lysis due to premature splitting, while high levels of CxaR, on the other hand, significantly delay cell splitting, leading to cell clumping and cells with multiple septa, by reducing Sle1 abundances.

*cxaR* was uncharacterized prior to this study, but its transcription has been shown to increase under various stress conditions, including treatment with vancomycin [71], oxacillin [72], mersacidin [73], brilacidin [74], and under heat shock [75]. This pattern of transcriptional upregulation in response to envelope damage is consistent with the finding that expression of *cxaR* likely is regulated by the two-component system VraSR (<u>v</u>ancomycin-resistance <u>a</u>ssociated <u>s</u>ensor/regulator). VraSR is activated by cell wall targeting antimicrobials, such as β-lactams and glycopeptides, and controls transcriptional induction of several cell wall-associated genes [72, 76]. While Sle1 and other PGHs are not directly regulated by the VraSR system, it can thus be speculated that VraSR-mediated upregulation of CxaR serves as a regulatory mechanism to reduce Sle1 levels under these conditions, thereby delaying cell splitting in response to cell wall stress. This delay could provide the cells with enough time to repair damaged peptidoglycan and reinforce cell wall integrity, ultimately enhancing bacterial survival.

While our results show that CxaR regulates Sle1 levels via ClpXP-mediated degradation, it may have additional roles in the cell that remain to be elucidated. For example, when performing localization microscopy with fluorescently tagged versions of CxaR, we surprisingly observed that overexpression of CxaR-GFP in Δ*clpX* appeared to improve growth of the deletion strain (**Fig 7E**). This was also confirmed with a non-tagged CxaR in the *clpX* deletion mutant (**S14 Fig**). Western blot assays did not indicate any decrease in cell wall-associated Sle1 during CxaR overexpression in Δ*clpX* cells (**S9 Fig**), which suggests that CxaR may have an additional role which helps the cells mitigate the impact of elevated Sle1 levels, potentially by rendering Sle1 non-functional. Additionally, the CxaR overexpression phenotype appears to be more severe than the expected defects from *sle1* deletion alone [2, 7, 9, 10], more closely resembling mutants with deletions of several PGHs [9, 77]. This could indicate an additional role of CxaR in cell division, or that CxaR facilitates ClpXP-mediated degradation of other proteins in addition to Sle1.

Sublethal concentrations of different antibiotics could rescue the growth defects observed in *cxaR* mutants, a phenomenon previously reported in *clpX* mutants [41]. For some antibiotics, the growth rescue can be directly attributed to reduced levels of Sle1 in the treated cells (**S10B-C Fig**). This is in line with previous observations that expression of certain PGHs, including *sle1,* is positively correlated with PBP transpeptidase activity [55, 56]. The reduction in Sle1 levels upon beta-lactam exposure is independent of *mecA*, as the same occurs in both MRSA (JE2) and non-MRSA (NCTC8325-4) strains [7]. Sle1 levels are also highly temperature dependent, increasing at 30°C and decreasing at temperatures exceeding 40°C [20]. The temperature-dependent growth phenotype of cells lacking CxaR is likely attributed to these fluctuating Sle1 levels. The exact mechanism by which temperature influences the levels of Sle1 is unknown, but the observed upregulation of CxaR under heat [75] and envelope stress [72, 76] suggest that elevated temperatures may enhance CxaR abundances, leading to increased Sle1 degradation.

Beyond the direct impact antibiotics have on hydrolase activity, additional mechanisms are likely involved in the antibiotic-mediated growth rescue of *cxaR* mutants. The growth stimulation mediated by sublethal concentrations of tunicamycin may be attributed to the role of teichoic acids in directing septal hydrolysis, in addition to its effect on Sle1 levels. A decrease in wall teichoic acid abundance has been shown to reduce septum-directed localization of PGHs and thus delay septal resolution [11, 15, 78, 79]. The same growth-rescue by tunicamycin has also been observed in a Δ*clpX* mutant [41], which additionally acquired spontaneous, growth-rescuing loss-of-function mutations in the gene encoding the lipoteichoic acid synthase, *ltaS* [42]. Furthermore, sublethal concentrations of β-lactams have been observed to prolong the duration of peptidoglycan biosynthesis at the septal wall, leading to cells with thickened, unresolved septa, due to the retention of septal PBPs [41, 53, 54]. A thicker septum could potentially mitigate the effects of increased autolytic activity caused by Sle1 accumulation in *cxaR* mutants.

*S. aureus* employs a complex network of regulatory mechanisms to effectively control the autolytic activity driving cell splitting. Given the important role of Sle1 in septum resolution, our findings establish CxaR as a key regulatory factor in maintaining the balance between peptidoglycan synthesis and hydrolysis – a balance which is critical to the function of antibiotics. As CxaR represents a previously unrecognized component in this process, further studies investigating its potential additional roles, as well as functional analogs in other species, may provide new insight into the mode of action of widely used antibiotic classes, including β-lactams.

## Materials and methods

### Bacterial strains and growth conditions

Strains used in this work are listed in **S3 Table**. All *S. aureus* strains were grown in tryptic soy broth (TSB) and *E. coli* strains were grown in lysogeny broth (LB) aerobically at 37°C, unless stated otherwise. Growth in liquid media was carried out in an orbital shaker at 180 rpm, while for growth on solid media 1.5% (w/v) agar was added to the respective medium base. When appropriate, the following antibiotics were added for selection: ampicillin (Amp, 100 µg/mL), chloramphenicol (Cam, 10 or 25 µg/mL), spectinomycin (Spc, 100 or 1,000 µg/mL for NCTC8325-4 and JE2, respectively), and erythromycin (Ery, 5 µg/mL). In addition, isopropyl β-D-1-thiogalactopyranoside (IPTG) or anhydrotetracycline (aTc) was added for induction of gene expression when needed.

For *E. coli* transformation, chemically competent IM08B cells were prepared using calcium chloride treatment, followed by heat shock transformation according to standard protocols. Plasmids isolated from *E. coli* IM08B were transformed into electrocompetent *S. aureus* by electroporation, as previously described [80].

### Strain and plasmid construction

See **S3, S4 and S5 Tables** for strains, plasmids, and oligonucleotides used in this work. Every construct was verified by PCR and sequencing.

#### Construction of CRISPRi knockdown strains

For gene knockdown, we used the two-plasmid CRISPR interference (CRISPRi) system described previously [59, 81, 82]. In this system, dCas9 is expressed from an IPTG-inducible promoter on the plasmid pLOW-*dcas9*, while the gene-specific single guide RNA (sgRNA) is constitutively expressed from a second plasmid (pCG248-sgRNA(*x*) or pVL2336-sgRNA(*x*), where *x* denotes the targeted gene). Upon co-expression, the dCas9-sgRNA complex binds the gene of interest thereby blocking transcription by the RNA polymerase. The sgRNA plasmids were constructed using inverse PCR for pCG248 [81] or Golden Gate cloning for pVL2336 [59], with the oligonucleotide primers listed in **S5 Table**.

#### Construction of the cxaR deletion mutant

*cxaR* was replaced by a spectinomycin resistance cassette in its native loci in *S. aureus* NCTC8325-4 using the temperature-sensitive pMAD system. First, the pMAD-*cxaR*::*spc* plasmid was constructed by assembling a fragment containing the *cxaR* up- and downstream regions (amplified using primer pairs mk591/mk592 and mk593/mk594, respectively) flanking a spectinomycin resistance cassette (amplified from pCN55 with primers mk188 and mk189) using overlap extension PCR. The fragment was ligated into pMAD using restriction cloning with NcoI and BamHI, and the resulting plasmid was transformed into *E. coli* IM08B. Construction of Δ*cxaR*::*spc* in strain NCTC8325-4 was then performed using a standard pMAD protocol with temperature shifts, as described before [83].

#### Construction of suppressor mutants

When the NCTC8325-4 *cxaR* deletion mutant was plated at 30°C, larger colonies consistently appeared among the very small colonies observed at 37°C. To determine whether these larger colonies had restored growth due to acquisition of spontaneous suppressor mutations, we performed a genetic suppressor screen similar to what has been described previously [42]. The Δ*cxaR* mutant was streaked for single colonies on TSA plates and incubated at 37°C overnight. Single small colonies were picked, resuspended in TSB, and grown overnight at 37°C with shaking. The following day, each independent overnight culture was plated on TSA and incubated overnight at 30°C. On some plates, visibly larger colonies had appeared. In those cases, a single large colony from each independent culture was selected, grown overnight in TSB at 30°C with shaking, and stored. For cultures without initially detectable larger colonies, a random colony was picked, grown overnight in TSB at 30°C, and then replated on TSA at 30°C. By the next day, all plates had a majority of large colonies. One large colony per plate was picked, grown overnight in TSB at 30°C, and stored. The growth of the stored cultures was analyzed, and five were selected for whole-genome re-sequencing.

#### Construction of plasmids for complementation and overexpression

*cxaR* and *clpX* were amplified from *S. aureus* genomic DNA using the primer pairs mhu28/mhu27 and mdb98/mdb99, respectively. The resulting fragments were digested with SalI and EcoRI and subsequently ligated into the corresponding restriction sites of plasmid pLOW-*m(sf)gfp*-*SA1477* to produce pLOW-*cxaR* and pLOW-*clpX*, which enable IPTG-inducible expression of the respective genes.

#### Construction of plasmids for localization analysis of CxaR and ClpX

To study the subcellular localization of C-terminally tagged CxaR, *cxaR* was amplified from *S. aureus* genomic DNA using primers mhu28 and mhu29, digested with SalI and BamHI, and ligated into the respective sites of pLOW-*SA0948*-*m(sf)gfp*. For N-terminally tagged CxaR, *cxaR* was amplified from *S. aureus* genomic DNA using primers mhu26 and mhu27, digested with BamHI and EcoRI, and ligated into the respective sites of pLOW-*m(sf)gfp*-*SA1477*. To study the subcellular localization of C-terminally tagged ClpX, *clpX* was amplified from *S. aureus* genomic DNA using primers mdb98 and mdb102, digested with SalI and BamHI, and ligated into the respective sites of pLOW-*lacA*-*mCherry*.

#### Construction of a chromosomally integrated cxaR-m(sf)gfp fusion

The temperature-sensitive pMAD system was used to GFP-tag *cxaR* in its native loci in *S. aureus* NCTC8325-4. To construct pMAD-*cxaR*-*m(sf)gfp*_*spc* the following four fragments were amplified: (1) the upstream region of *cxaR* with primers mdb58 and mdb59 using gDNA from *S. aureus* NCTC8325-4 as a template, (2) the *cxaR*-*m(sf)gfp* fusion from plasmid pLOW-*cxaR*-*m(sf)gfp* using primers mdb60 and mdb61, (3) the spectinomycin resistance cassette using primers mk503 and mk504 from plasmid pCN55, and (4) the downstream region of *cxaR* from *S. aureus* NCTC8325-4 gDNA using mdb62 and mdb63. These fragments were subsequently fused by overlap extension PCR and ligated into pMAD using the NcoI and SalI restriction sites introduced by the outer primers. Finally, chromosomal integration of the fusion construct was performed following a standard pMAD protocol [84].

#### Construction of split luciferase plasmids

To C-terminally fuse CxaR, ClpX, and ClpX_I265E_ to the small luciferase subunit (SmBit), *cxaR*, *clpX*, and *clpX_I265E_* were amplified from *S. aureus* NCTC8325-4 WT or *clpX_I265E_* genomic DNA using the primer pairs efs7/efs8 for *cxaR* and mdb93/mdb94 for *clpX* and *clpX_I265E_*. The amplified fragments and vectors pAF256-P_tet_-*hupA-smbit*/*lgbit* and pAP118-P_tet_-*hupA-smbit*/*hupA-lgbit* were then digested with SacI and XhoI. The digested fragments were ligated into the vectors and subsequently transformed into *E. coli* IM08B. The pAF256 vectors, encoding CxaR, ClpX, or ClpX_I265E_ fused to SmBit and LgBit not part of a fusion protein, served as negative controls in the split luciferase assays. The pAP118 vectors were further modified to C-terminally fuse CxaR, ClpX, ClpP, Sle1, or Sle1 (without signal peptide) to the large luciferase subunit (LgBit). All fragments were amplified using NCTC8325-4 WT genomic DNA as template and the following primer pairs: (1) efs20/efs21 for *cxaR*, (2) efs5/efs6 for *clpX*, (3) efs9/efs10 for *clpP*, (4) mdb88/mdb89 for *sle1*, and (5) mdb90/mdb89 for *sle1* (without signal peptide). The amplified fragments and the corresponding pAP118 vectors were then digested with PvuI and NotI, ligated, and subsequently transformed into *E. coli*.

### Genome re-sequencing and analysis

Whole-genome sequencing of Δ*cxaR* (MK2047), the strain expressing chromosomally integrated CxaR-GFP (MDB376), and the *cxaR* suppressor mutants (EFS104, EFS105, MDB390, MDB401, and MDB404) was performed. Genomic DNA was isolated from the respective strains using The Wizard Genomic DNA Purification Kit (Promega) and sent to Novogene for Illumina sequencing. Sequence assembly to the reference genomes and SNP detection were done using Geneious Prime [85].

### CRISPRi-sequencing screen

The explorative CRISPRi-seq screen for penicillin G susceptibility determinants was performed as described before, using an already constructed genome-wide CRISPRi library in the *S. aureus* strain NCTC8325-4 [59, 60]. Briefly, the library (pre-grown in BHI to OD_600_ = 0.8) was diluted 1:1,000 in 100 mL ultra-high temperature (UHT) treated milk (Tine SA, Norway) and incubated at 37 °C with shaking for 7 hours. After, it was reinoculated 1:1,000 in fresh UHT milk and grown for another 6 hours. Chloramphenicol (10 µg/mL) was added for selection, and when appropriate, 30 ng/mL anhydrotetracycline was added for induction of the CRISPRi- system. Furthermore, to identify fitness determinant affected by β-lactam treatment, the library was grown in the presence and absence of 0.008 µg/mL penicillin G (corresponding to the MIC for NCTC8325-4 in milk). Four parallels were included per condition. After growth, cells were harvested by centrifugation, and plasmids were isolated as previously described [60]. Illumina amplicon libraries were prepared, and sequencing was performed on a NovaSeq platform using 150 bp paired-end reads (performed by Novogene, UK) following established protocols [60]. CRISPRi-seq differential enrichment analyses was done as described by Bakker et al. [58] and Mårli et al. [60]. Notably, due to the inhibitory concentration of penicillin G used in this explorative screen, the number of generations for growth of the penicillin G treated library was significantly lower than for the untreated library.

### Growth assays in liquid media

To assess bacterial growth in a liquid medium, overnight cultures of the strains to be monitored were diluted 1:1,000 in TSB medium supplemented with the appropriate antibiotics and inducers. For MIC analysis, the following panel of antibiotics were tested against the JE2 *cxaR* knockdown mutant and CRISPRi control strain in two-fold dilution series: penicillin G (PEN), oxacillin (OXA), cefoxitin (CXI), vancomycin (VAN), teicoplanin (TEC), bacitracin (BAC), moenomycin (MOE), tunicamycin (TUN), ciprofloxacin (CIP), novobiocin (NOV), trimethoprim (TMP), rifampicin (RIF), erythromycin (ERY), kanamycin (KAN), and streptomycin (STR). Bacterial dilutions were added to a 96-well microtiter plate and incubated in a plate reader at 37°C for 20 hours, unless otherwise stated. OD_600_ measurements were taken every 10 minutes, with a brief shaking of the plate for 5 seconds prior to each measurement. All growth curves in this work are the mean of three replicate measurements, with the exception of the MIC panel (**S3 Fig**), growth at different temperatures (**S6A Fig**), and CxaR/ClpX overexpression (**S14 Fig**) which represent the mean value of two replicates, and they are all representative of at least two independent experiments.

### Spotting assays

Overnight cultures diluted 1:1,000 in TSB with the appropriate antibiotics were grown at 37°C until they reached an OD_600_ of 0.5 or the indicated time point. The cultures were diluted in a ten-fold dilution series from 10^−1^ to 10^−8^ and 5 µL from each dilution was spotted on TSA. The TSA, in addition to the appropriate selective antibiotics, contained 500 µM IPTG for CRISPRi knockdown strains and 1 ng/mL penicillin G or 6 ng/mL oxacillin when assessing growth rescue by β-lactams. The plates were incubated for 24 hours at their indicated temperature. All presented spotting assays are representative of three technical replicates from at least three independent experiments.

### Western blot analyses of Sle1 levels

#### Cell harvesting

Overnight cultures were diluted 1:1,000 in TSB medium with the appropriate antibiotics and inducers and incubated at either 37°C or 42°C with shaking until an OD_600_ of approximately 0.4 was reached.

#### Preparation of extracellular and cell wall-associated protein extracts

Culture volumes of 50 mL were harvested by centrifugation at 6,000 x *g* for five minutes at 4°C. To extract extracellular proteins, 30 mL of the supernatants were mixed with 30 mL 96% ethanol and incubated overnight at 4°C. To extract cell wall-associated proteins, the pellets were washed with 10 mL cold 0.9% (w/v) NaCl and centrifuged at 6,000 x *g* at 4°C for five minutes. The supernatants were discarded, and cell wall-associated proteins were eluted with 10 mL 4% (w/v) SDS at 25°C for 45 minutes with shaking. The samples were then centrifuged at 6,000 x *g* for five minutes at 4°C, and the resulting supernatants were mixed with 10 mL 96% ethanol and left to precipitate at 4°C overnight. The following day, the protein extracts were pelleted by centrifugation at 10,000 x *g* for 30 minutes at 4°C, dried in a fume hood, and normalized with 50 mM Tris-HCl (pH 7.4). The normalized extracts were mixed 1:1 with 2x SDS loading buffer and heated at 95°C for five minutes prior to SDS-PAGE.

#### Preparation of whole cell lysate

Culture volumes of 50 mL, normalized to an OD_600_ of 0.4, were harvested by centrifugation at 4,000 x *g* for five minutes at 4°C. The pellets were resuspended in 100 µL TBS buffer and mechanically lysed using the Fast Prep method with ≤106 µm glass beads in three cycles of 20 seconds at 6 m/s, with one minute incubation on ice between cycles. Insoluble material was removed by centrifugation at 20,000 × g for 2 minutes. Next, the supernatants were mixed with equal volumes 2x SDS loading buffer and heated at 95°C for 5 minutes prior to SDS-PAGE.

#### SDS-PAGE, Western blotting, and detection

The protein samples were separated by SDS-PAGE on a polyacrylamide gel, which consisted of a 12% separation gel with a 4% stacking gel layered on top. The proteins were then electroblotted onto a PVDF membrane using the Trans-Blot Turbo System (Bio-Rad). Afterward, the membrane was trimmed to retain proteins below 60 kDa and blocked in TBST, containing 5% (w/v) bovine serum albumin (BSA) and 1 μg/mL human IgG, for 1 hour at room temperature, followed by overnight incubation at 4°C. After washing the membrane with TBST, the following day, it was incubated with an anti-Sle1 primary antibody [20] (diluted 1:10,000 in TBST with 2% (w/v) BSA and 1 μg/mL human IgG) for 1 hour. Next, unbound antibodies were removed by three TBST washes, and the membrane was then incubated for another hour with an anti-mouse IgG HRP-conjugate secondary antibody (Promega) (diluted 1:10,000 in TBST with 1 μg/mL human IgG). After another series of washes, the membrane was finally developed using the SuperSignal West Pico PLUS Chemiluminescent Substrate kit (Thermo Fisher Scientific), and blot images were captured with an Azure Imager c400 (Azure Biosystems).

### Western blot analysis of GFP-tagged CxaR

Overnight cultures of MDB376 (NCTC8325-4 with a chromosomally integrated *cxaR-m(sf)gfp* fusion), MDB430 (NCTC8325-4 carrying pLOW-*cxaR-m(sf)gfp*), and MDB395 (NCTC8325-4 *clpX*_I265E_ carrying pLOW-*cxaR-m(sf)gfp*) were diluted 1:1,000 in TSB medium supplemented with 100 µg/mL spectinomycin (for MDB376) or 5 µg/mL erythromycin and 10 µM IPTG (for MDB430 and MDB395). The bacterial cultures were incubated at 37°C until they reached an OD_600_ of approximately 0.5 and 1. They were then normalized to an OD_600_ of 0.5, and harvested by centrifugation at 4,000 × *g* for 5 minutes at 4°C. The pellets were resuspended in 200 µL TBS buffer and lysed mechanically using the Fast Prep method with ≤106 µm glass beads at 6 m/s for three cycles of 20 seconds each. Insoluble material was removed by centrifugation at 20,000 × g for 2 minutes. Next, the supernatants were mixed with equal volumes 2x SDS loading buffer and heated at 95°C for 5 minutes.

The subsequent SDS-PAGE, blotting, and GFP detection were performed as described above for Sle1 immunoblotting, with the exception of antibody selection. GFP-tagged CxaR was detected using an anti-GFP primary antibody (Invitrogen) (diluted 1:4,000 in TBST) and an anti-rabbit IgG HRP-conjugated secondary antibody (Promega) (diluted 1:5,000 in TBST).

### Phase contrast and epifluorescence microscopy

#### Microscopy

Overnight cultures were diluted 1:1,000 in TSB medium with the appropriate antibiotics and inducers. These cultures were incubated at 37°C with shaking until they reached the desired optical density (OD_600_ ∼0.4) or time point, followed by cooling on ice. In some cases, cells were then stained with fluorescent vancomycin (VanFL, 0.8 µg/mL [Invitrogen]) to visualize the cell wall and/or 4′,6-diamidino-2-phenylindole (DAPI, 7.5 µg/mL [Invitrogen]) to visualize the nucleoid. Bacterial cells were immobilized on 1.2% (w/v) agarose pads before imaging on either a Zeiss Axio Observer microscope or a Leica DMi8 microscope. The bacteria were visualized with a 100× phase contrast objective, and images were captured using an ORCA-Flash4.0 V3 Digital CMOS (for the Zeiss microscope) or a K8 Scientific CMOS (for the Leica microscope) camera.

#### Image analysis

Cell size distribution among different *S. aureus* strains was determined using MicrobeJ [86]. Bacterial cells were automatically detected using phase contrast image stacks, and every image was subsequently manually corrected by adding unidentified cells or removing incorrectly identified cells. Additionally, co-localization of CxaR-GFP and ClpX-mCherry was analyzed manually by examining both signals in 300–350 randomly selected cells from two biological replicates (n = 635).

### Transmission electron microscopy

Overnight cultures of the NCTC8325-4 *cxaR* CRISPRi knockdown and control strains were diluted 1:1,000 in 10 mL TSB with the appropriate selective antibiotics and inducers, with and without 1 ng/mL penicillin G. The cultures were grown at 37°C for 11 hours, corresponding to the transient minimum of the *cxaR* knockdown strain. The cultures were added to fixative to a final concentration of 1.25% (v/v) glutaraldehyde, 2% (v/v) paraformaldehyde, and 100 mM cacodylate (CaCo) buffer. The cells were fixed for one hour at room temperature and overnight at 4°C, before being washed in 100 mM CaCo buffer three times. The final pellets were embedded in 3% (w/v) low-melt agarose and stored in 100 mM CaCo buffer. Post-fixation with 1% osmium tetroxide was performed for one hour. Excess osmium tetroxide was removed by washing with 100 mM CaCo buffer and the pellets were dehydrated using gradually increasing concentrations of cold acetone. The dehydrated pellets were embedded in EPON epoxy resin and polymerized at 60°C. The samples were sectioned 60 nm thick and stained with uranyl acetate and lead citrate. A Jeol JEM-2100Plus was used for imaging.

### Split luciferase assays

Pairwise protein-protein interactions were assessed *in vivo* in *S. aures* NCTC8325-4 using the split luciferase system developed by Paiva et al. [68] for *C. difficile*, following the same approach as described before [87].

### Structural and phylogenetic analysis

The presence of general structural features, including transmembrane helices and signal peptides, were predicted using SignalP-6.0 and DeepTMHMM [88, 89]. Homologs of CxaR within *Staphylococcaceae* were identified through BLASTp, using the NCTC8325-4 CxaR peptide sequence as the query [90]. Sequence representatives from 55 species within *Staphylococcus*, *Mammaliicoccus*, and *Macrococcus* were aligned by Clustal Omega, and the multiple sequence alignment (MSA) was visualized using Jalview [91, 92]. Phylogenetic trees from the homologs were generated using MEGA11 [93]. HMMER and HHpred were applied for sensitive hidden Markov model-based homolog searches to detect structural motifs [61, 62]. TPRpred from the HH-suite was used to detect TPR-like motifs in CxaR homologs [94].

### *In silico* protein interaction modelling

AlphaFold 3 [95] was used for the modelling of every protein structure and protein interactions in this work. The top ranked predictions were selected and visualized using either ChimeraX [96] or PyMOL [97].

## Acknowledgements

We would like to thank the NMBU Imaging Center for help with TEM, and Zhian Salehian and Anne Ohren Nordraak at NMBU for laboratory assistance.

