## Supplementary material for "Cell splitting in *Staphylococcus aureus* is controlled by an adaptor protein facilitating degradation of a peptidoglycan hydrolase": Supplmental information

**This PDF file includes:**

S1 to S14 Figures

Legend for S1 Table

S2 to S5 Tables

SI References

**Other supporting materials for this manuscript include the following:**

S1 Table

### Supporting figures

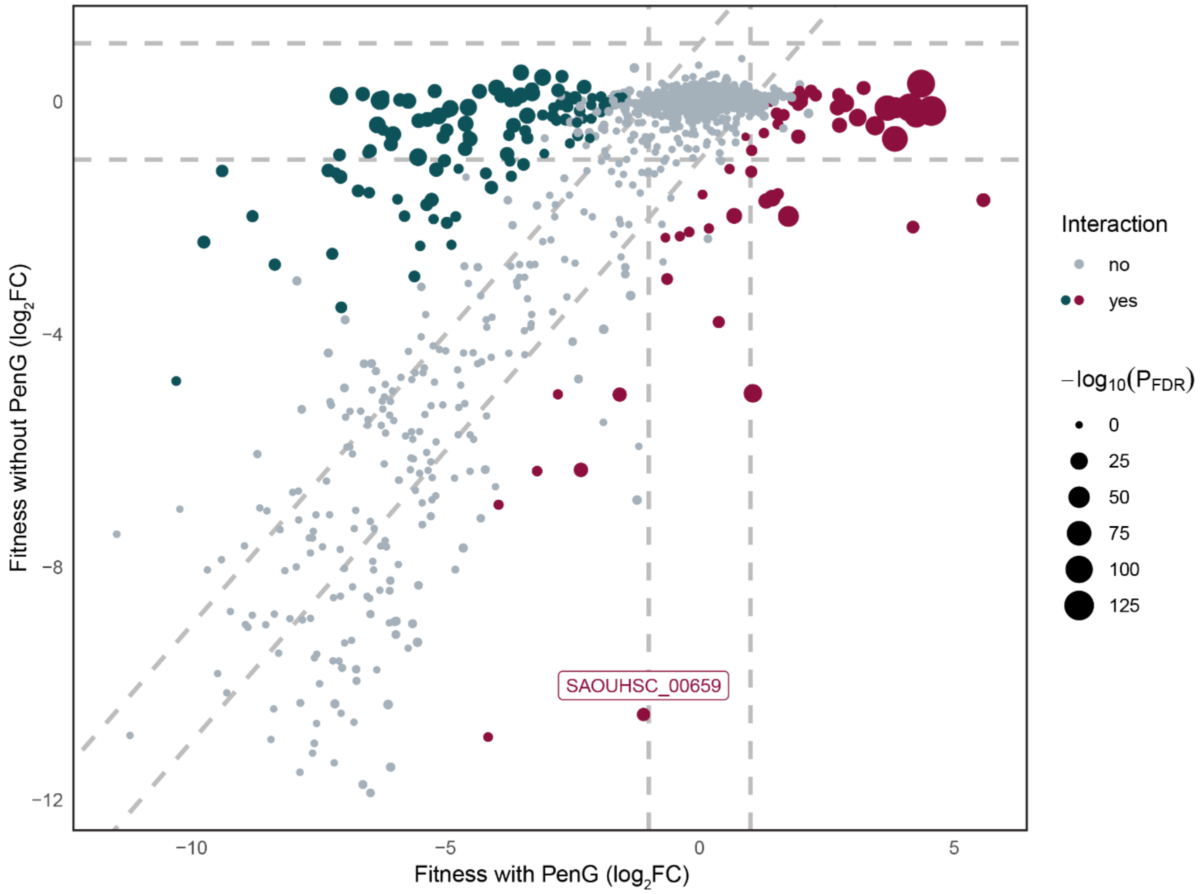


**S1 Fig. CRISPRi-seq identified genetic factors modulating susceptibility to β-lactams in *S. aureus*.** The growth fitness impact of knocking down each transcriptional unit was assessed under conditions with (*x-axis*) and without (*y-axis*) penicillin G (0.008 μg/ml). sgRNAs found to be significantly reduced upon penicillin G treatment, indicating that knockdown of these transcriptional units increases sensitivity to penicillin G, are highlighted in blue ($log_2FC < -1$, $P_{adj} < 0.05$). In contrast, sgRNAs that were significantly enriched, indicating that their knockdown reduces sensitivity to penicillin G, are highlighted in red ($log_2FC > 1$, $P_{adj} < 0.05$). Among the significantly enriched sgRNA, *SAOUHSC_00659* (*cxaR*) was the top hit.

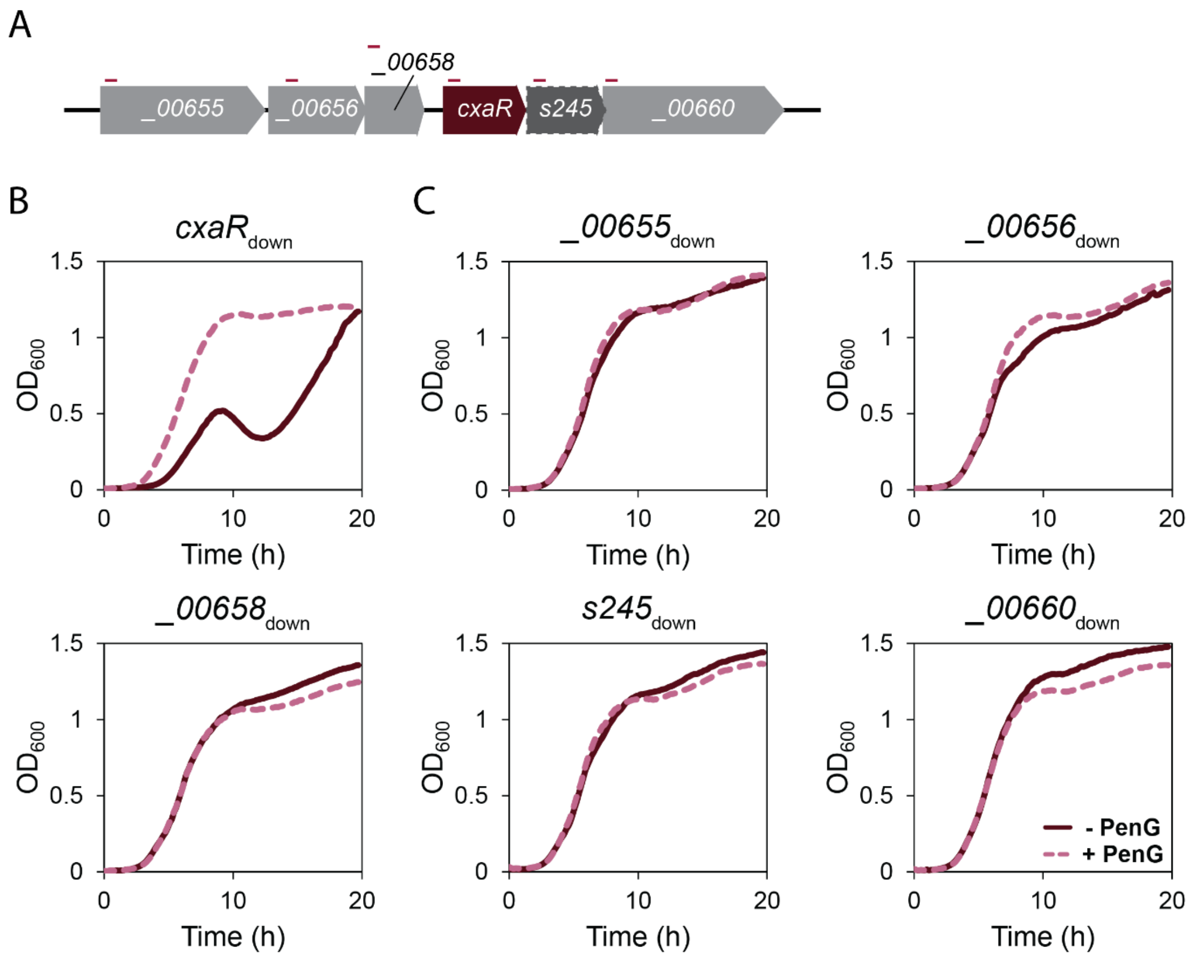


**S2 Fig. The phenotype observed upon knockdown or deletion of *cxaR* is not caused by polar effect on neighboring genes.** (**A**) Schematic overview of the genes surrounding *cxaR* in its operon. Red lines indicate the approximate location of sgRNA targets used for CRISPRi knockdown. (**B**) Growth curves of an NCTC8325-4 $cxaR_{down}$ strain in TSB at 37°C with (dashed line) and without (solid line) 1 ng/mL of penicillin G. Knockdown was induced by 30 ng/mL aTc. (**C**) Growth curves showing the effects of CRISPRi knockdown of genes in NCTC8325-4 located upstream (*SAOUHSC_00655*, *SAOUHSC_00656* and *SAOUHSC_00658*) and downstream (*S245* and *SAOUHSC_00660*) of *cxaR*, with (dashed line) and without (solid line) 1 ng/mL of penicillin G. The strains were grown at 37°C in TSB with 30 ng/mL aTc to induce knockdown.

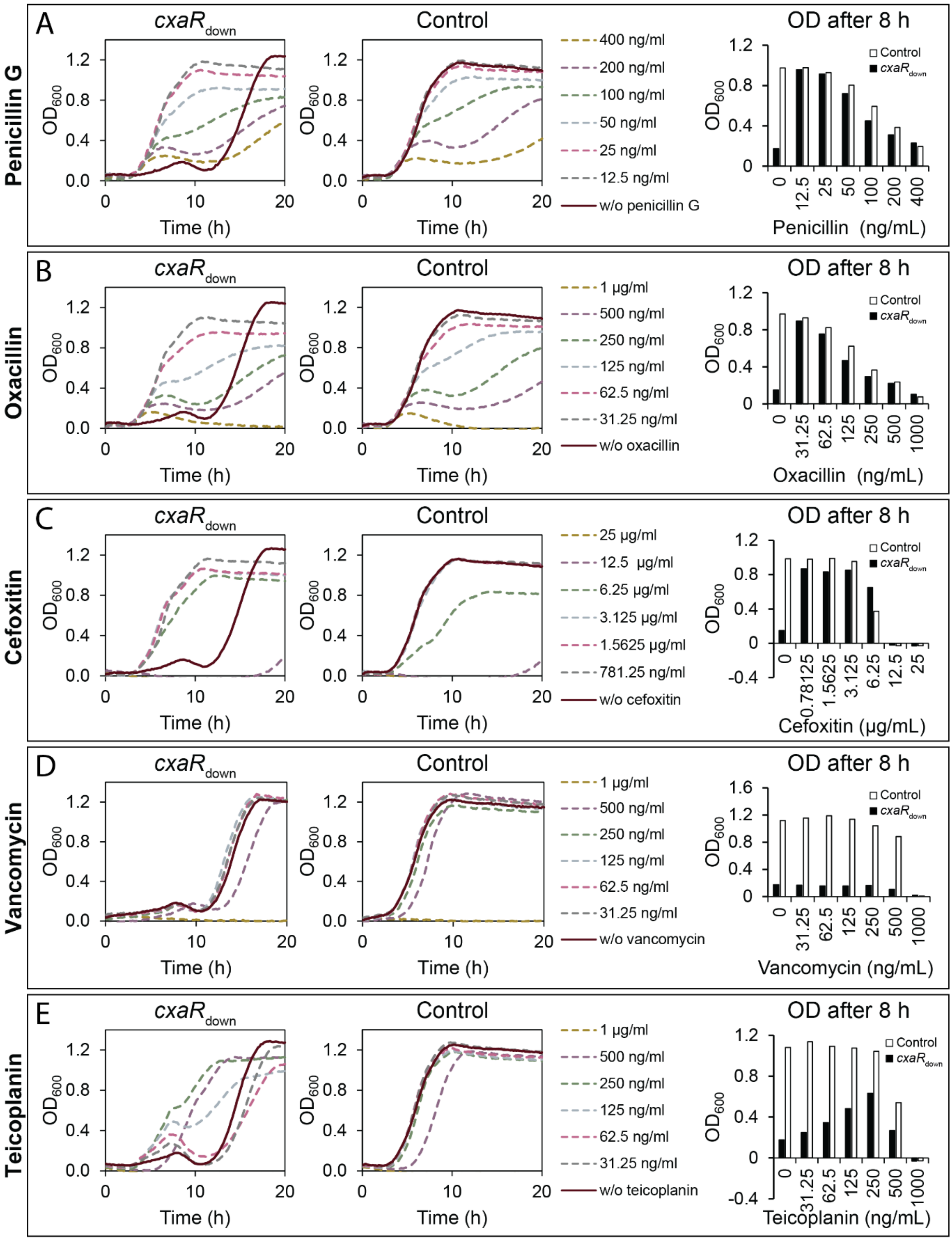
A
Penicillin G
cxaR down
Control
OD after 8 h
400 ng/ml
200 ng/ml
100 ng/ml
50 ng/ml
25 ng/ml
12.5 ng/ml
w/o penicillin G
Penicillin (ng/mL)
B
Oxacillin
1 µg/ml
500 ng/ml
250 ng/ml
125 ng/ml
62.5 ng/ml
31.25 ng/ml
w/o oxacillin
Oxacillin (ng/mL)
C
Cefoxitin
25 µg/ml
12.5 µg/ml
6.25 µg/ml
3.125 µg/ml
1.5625 µg/ml
781.25 ng/ml
w/o cefoxitin
Cefoxitin (µg/mL)
D
Vancomycin
1 µg/ml
500 ng/ml
250 ng/ml
125 ng/ml
62.5 ng/ml
31.25 ng/ml
w/o vancomycin
Vancomycin (ng/mL)
E
Teicoplanin
1 µg/ml
500 ng/ml
250 ng/ml
125 ng/ml
62.5 ng/ml
31.25 ng/ml
w/o teicoplanin
Teicoplanin (ng/mL)
OD600
Time (h)

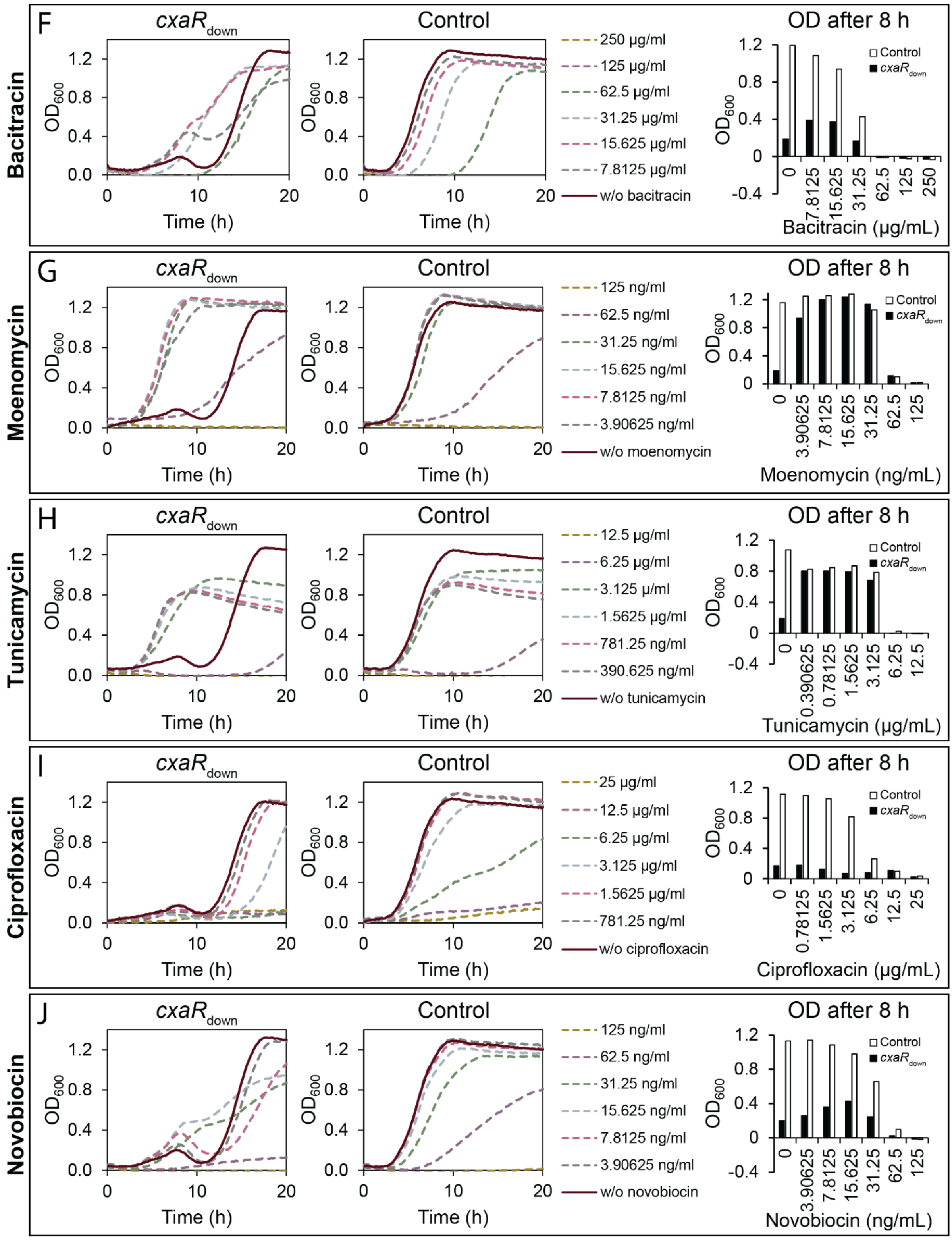

F
Bacitracin
cxaR_down
Control
OD after 8 h
250 µg/ml
125 µg/ml
62.5 µg/ml
31.25 µg/ml
15.625 µg/ml
7.8125 µg/ml
w/o bacitracin
Time (h)
OD600
Bacitracin (µg/mL)
G
Moenomycin
125 ng/ml
62.5 ng/ml
31.25 ng/ml
15.625 ng/ml
7.8125 ng/ml
3.90625 ng/ml
w/o moenomycin
Moenomycin (ng/mL)
H
Tunicamycin
12.5 µg/ml
6.25 µg/ml
3.125 µ/ml
1.5625 µg/ml
781.25 ng/ml
390.625 ng/ml
w/o tunicamycin
Tunicamycin (µg/mL)
I
Ciprofloxacin
25 µg/ml
12.5 µg/ml
6.25 µg/ml
3.125 µg/ml
1.5625 µg/ml
781.25 ng/ml
w/o ciprofloxacin
Ciprofloxacin (µg/mL)
J
Novobiocin
125 ng/ml
62.5 ng/ml
31.25 ng/ml
15.625 ng/ml
7.8125 ng/ml
3.90625 ng/ml
w/o novobiocin
Novobiocin (ng/mL)

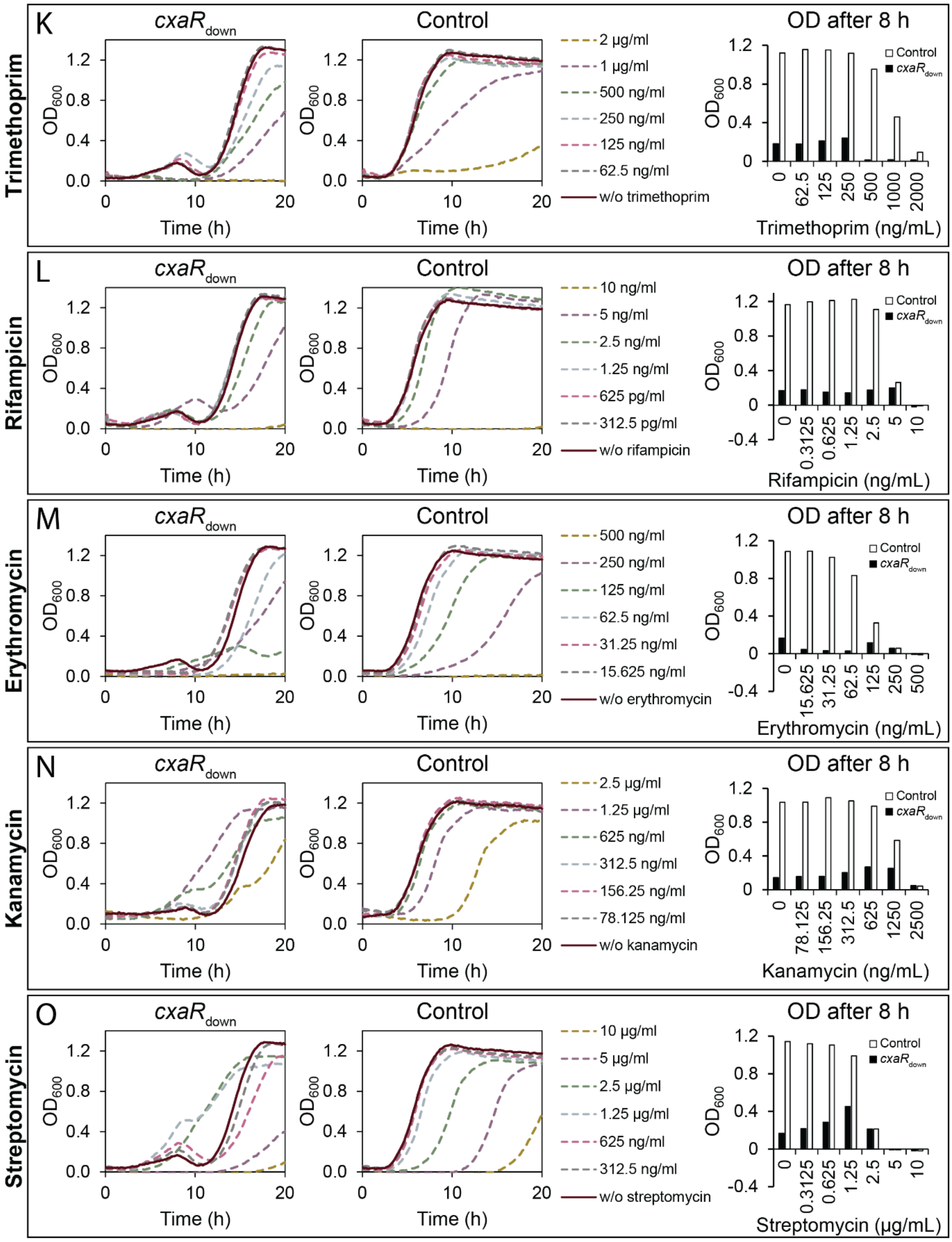


**S3 Fig. Sublethal concentrations of various antibiotics stimulate growth of cells lacking *cxaR*.** The JE2 *cxaR*$_{down}$ strain (left panels) and the CRISPRi control (middle panels) subjected to various concentrations of different antibiotics during growth in TSB at 37°C. Knockdown was induced by 500 µM IPTG. The antibiotics tested included the β-lactams (**A**) penicillin G, (**B**) oxacillin, and (**C**) cefoxitin, the glycopeptides (**D**) vancomycin and (**E**) teicoplanin, the polypeptide antibiotic (**F**) bacitracin, the phosphoglycolipid antibiotic (**G**) moenomycin, the WTA-inhibiting nucleoside antibiotic (**H**) tunicamycin, the fluoroquinolone (**I**) ciprofloxacin, the aminocoumarin (**J**) novobiocin, the folate biosynthesis inhibitor (**K**) trimethoprim, the ansamycin (**L**) rifampicin, the macrolide (**M**) erythromycin, and the aminoglycosides (**N**) kanamycin and (**O**) streptomycin. The right panels show the $OD_{600}$ for each strain after 8 hours of growth for each concentration of the antibiotics tested.

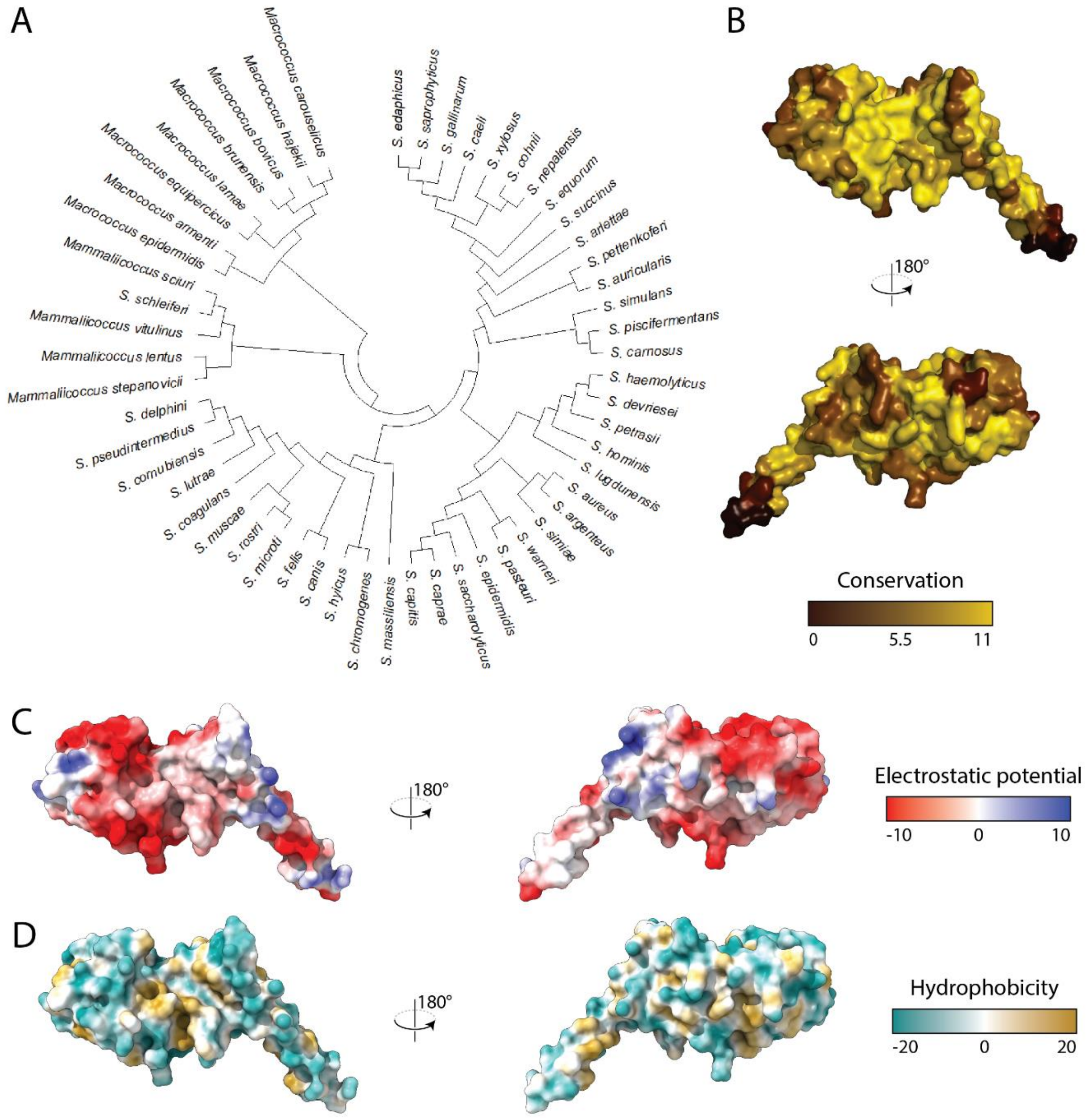


**S4 Fig. CxaR is conserved among *Staphylococcaceae*.** (**A**) Phylogenetic tree of CxaR homologues from 55 species within the *Staphylococcaceae* family. The phylogenetic relatedness of the homologues generally clustered by genera, with the exception of *Staphylococcus schleiferi* which appeared to utilize a CxaR homolog more closely resembling those found in *Mammaliicoccus*. (**B**) AlphaFold 3-predicted structure of CxaR with residue conservation from the phylogenetic analysis in **A** mapped on to it. The colors represent conservation scores ranging from 0 (least conserved) to 11 (most conserved), accounting for both conservation of specific amino acids and their stereochemical properties. (**C**) Coulombic electrostatic potential of the predicted CxaR structure, with red for negative potential through white to blue for positive potential. (**D**) Hydrophobicity (molecular lipophilicity potential) of the predicted CxaR structure, with dark cyan for most hydrophilic through white to dark goldenrod for most hydrophobic.

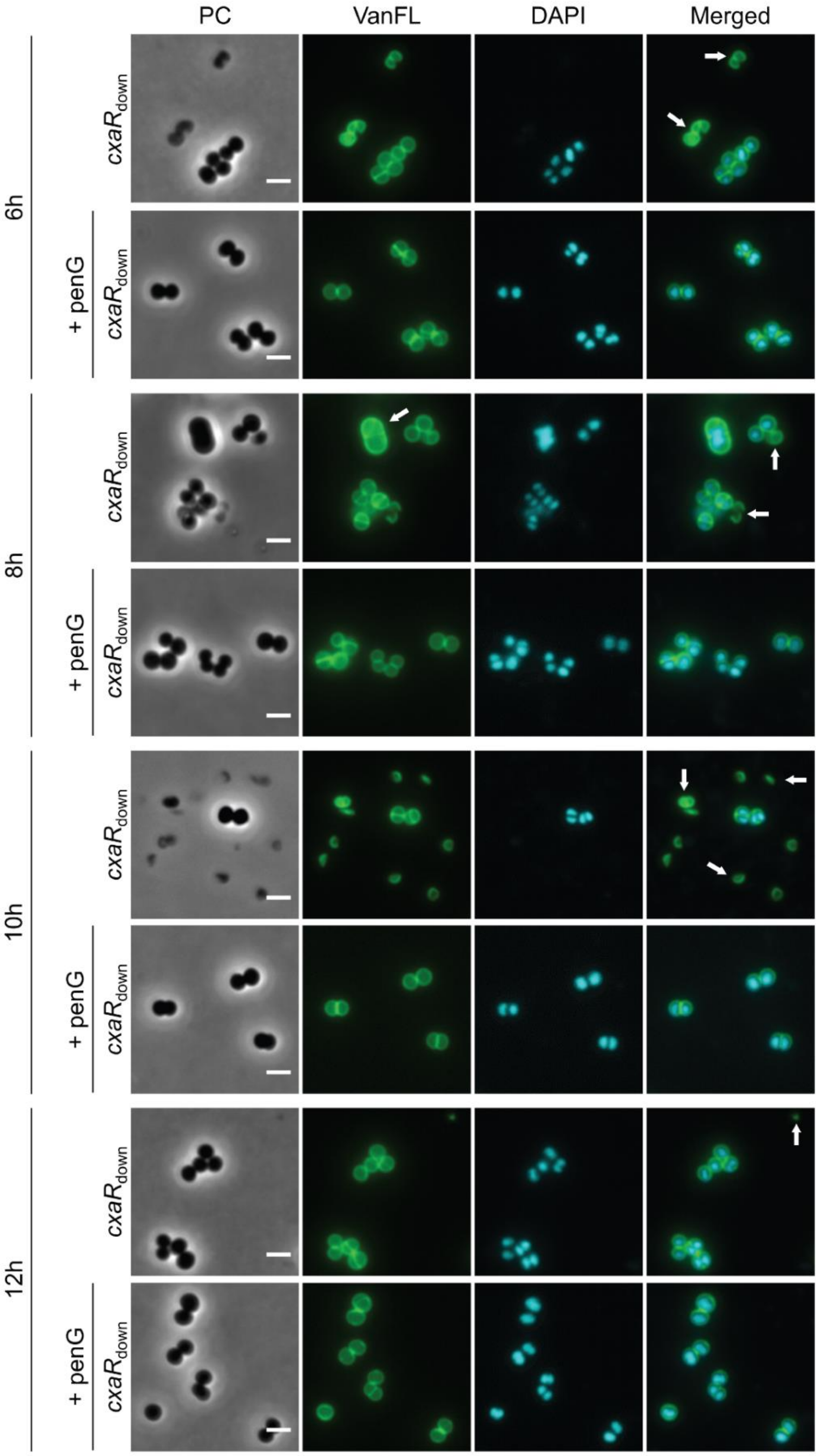


**S5 Fig. Sublethal concentrations penicillin G significantly reduce lysis of cell lacking *cxaR*.** Phase contrast and fluorescence microscopy of the JE2 $cxaR_{down}$ strain after 6, 8, 10, and 12 hours of incubation in TSB at 37°C, with and without 5 ng/mL penicillin G. CRISPRi knockdown of *cxaR* expression was induced by adding IPTG to a final concentration of 500 µM. The cells were stained with VanFL (cell wall) and DAPI (nucleoid). White arrows indicate abnormal and recently lysed cells. Scale bars represent 2 µm.

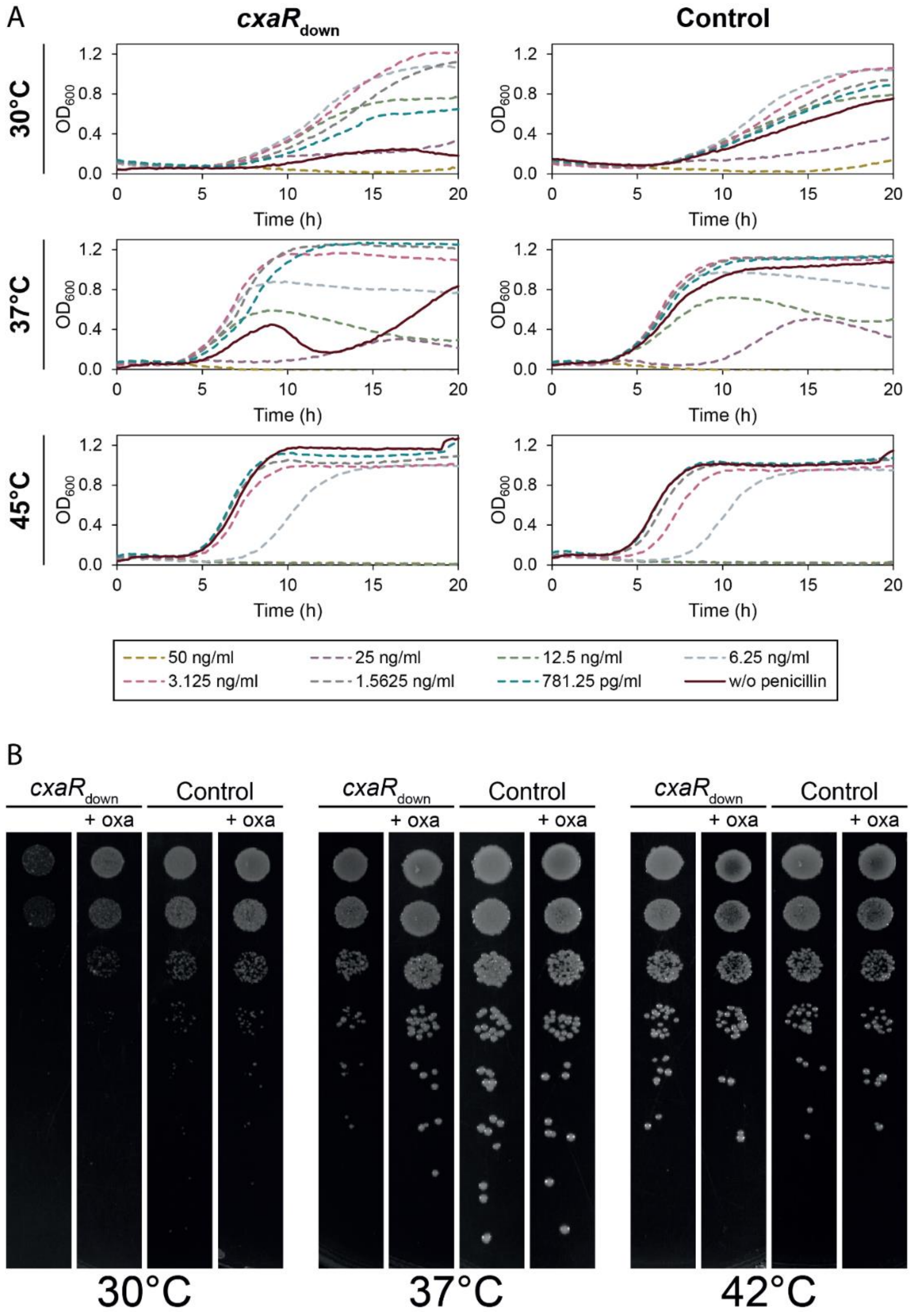


**S6 Fig. *cxaR* mutants exhibit heat stability and cold sensitivity.** (**A**) The NCTC8325-4 *cxaR*$_{down}$ and CRISPRi control strains grown in TSB at 30°C, 37°C, and 45°C with different concentrations of penicillin G. Gene knockdown was induced with the addition of 500 µM IPTG. The graphs represent averages from duplicate measurements. (**B**) Spotting of the NCTC8325-4 *cxaR*$_{down}$ and CRISPRi control strains at different temperatures with or without addition on oxacillin. The strains were grown until an $OD_{600}$ of 0.5 was reached, serially diluted in 10-fold dilution series, and 5 µL of each dilution was spotted on TSA containing 500 µM IPTG, with or without 6 ng/mL oxacillin. The plates were incubated at 30°C, 37°C, or 42°C for 24 hours before imaging.

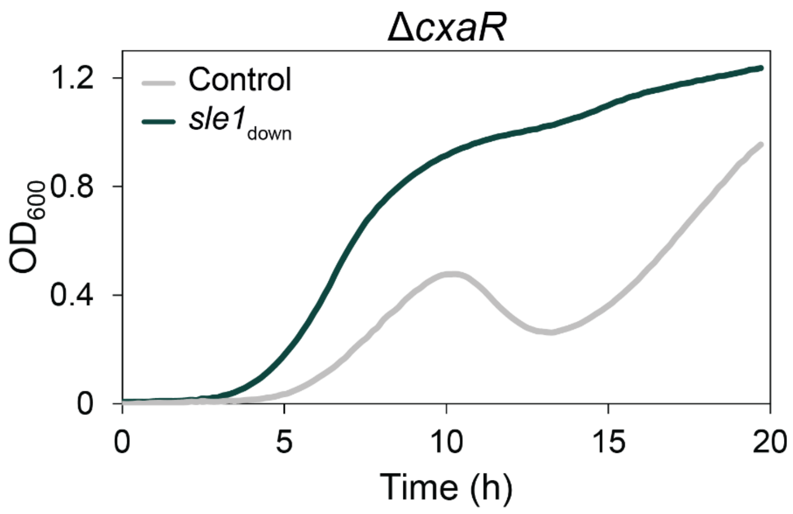


**S7 Fig. Reduced Sle1 activity can compensate for the lack of CxaR.** Growth of an NCTC8325-4 Δ*cxaR* mutant with (green) and without (grey) knockdown of *sle1* in TSB at 37°C. Gene knockdown was induced with the addition of 500 µM IPTG. The graphs represent averages from triplicate measurements.

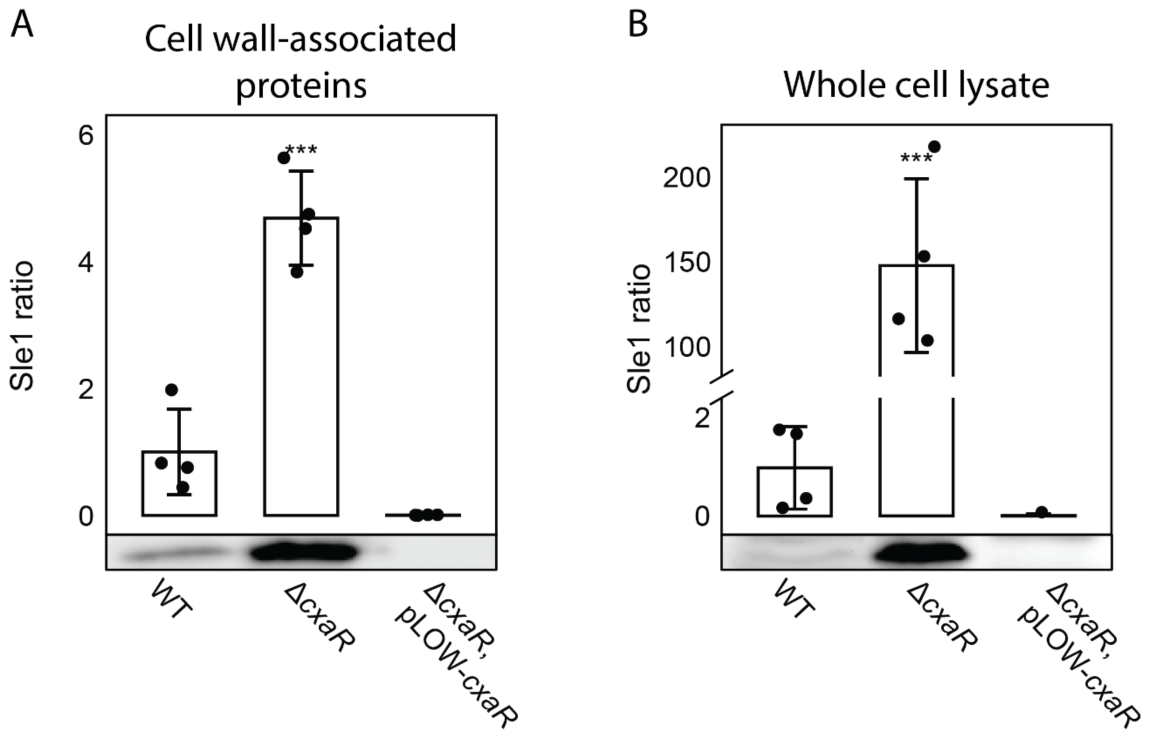


**S8 Fig. Sle1 levels are elevated in the absence of CxaR and reduced upon CxaR overexpression.** Western blot analysis of Sle1 levels in (**A**) cell wall-associated protein fractions and (**B**) whole-cell lysates from NCTC8325-4 wild-type, *cxaR* deletion, and *cxaR* overexpressing cells. Band intensities were quantified using ImageJ and normalized to the wild-type signal. (**A**) In cell wall-associated fractions, Sle1 levels were increased in Δ*cxaR* cells (4.681 ± 1.270) and reduced in CxaR overexpressing cells (0.007 ± 0.003), relative to the wild-type (1.000 ± 0.220). (**B**) Similarly, in whole-cell lysates, Sle1 levels were highly elevated in Δ*cxaR* cells (147.847 ± 45.768) and depleted in CxaR overexpressing cells (0.015 ± 0.043), compared to the wild-type (1.000 ± 1.095). Statistical significance was assessed using one-way ANOVA followed by Tukey's Honest Significant Difference (HSD) test (***, *P*-value < 0.001). Column height represents the mean, and error bars the standard deviation from four biological replicates. Representative Sle1 blots are shown below the corresponding graphs.

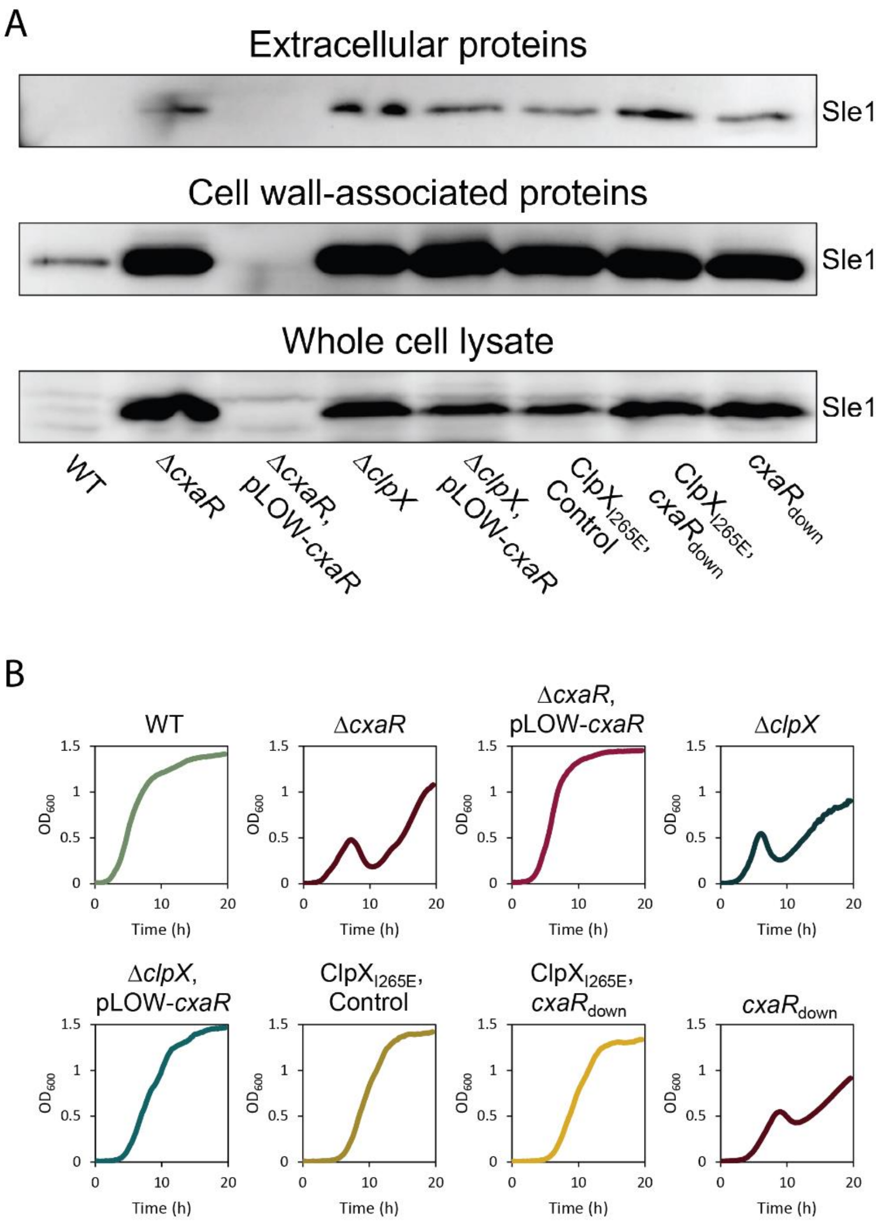


**S9 Fig. CxaR regulates Sle1 levels in a ClpXP-dependent manner.** (**A**) Western blots of extracellular, cell wall-associated, and whole cell Sle1 levels in the NCTC8325-4 wildtype, Δ*cxaR* mutant, *cxaR* overexpressing strain, Δ*clpX* mutant, Δ*clpX* mutant overexpressing *cxaR*, $clpX_{I265E}$ mutant, $clpX_{I265E}$ mutant with knockdown of *cxaR*, and *cxaR* knockdown strain. Gene knockdown and overexpression were induced with the addition of 500 µM IPTG. (**B**) Growth curves displaying the growth phenotypes of the strains Western blotted in **A** at 37°C in TSB. Gene knockdown and overexpression were induced with the addition of 500 µM IPTG. The graphs represent averages from triplicate measurements.

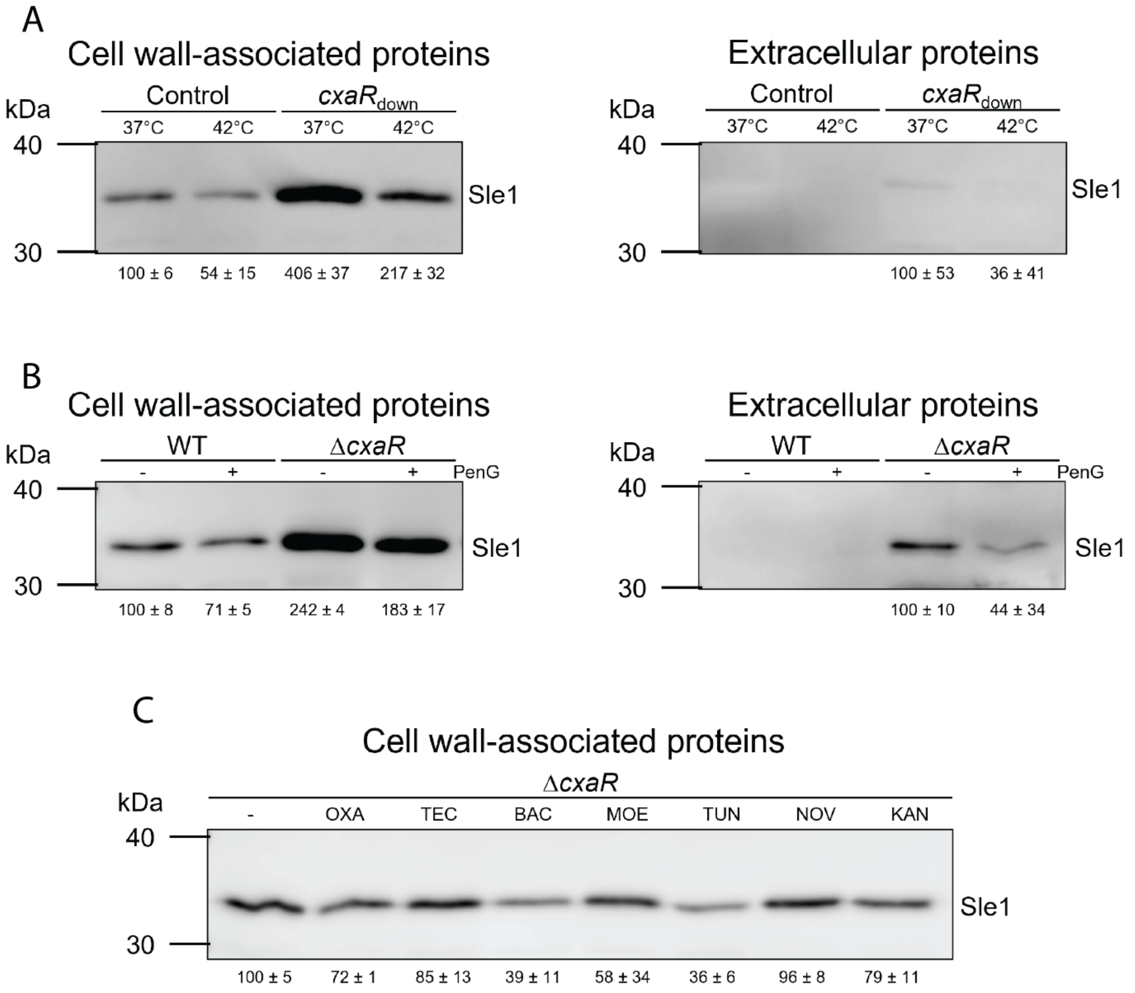


**S10 Fig. Supraoptimal temperatures and various antibiotics reduce the level of cell wall-associated Sle1.** (**A**) Western blots displaying the difference in cell wall-associated and extracellular Sle1 levels in the NCTC8325-4 *cxaR*$_{down}$ strain and CRISPRi control incubated at 37°C and 42°C. IPTG (500 µM) was added for gene knockdown. Band intensities were quantified using ImageJ. Values for cell wall-associated Sle1 were normalized to the control band at 37 °C, while extracellular Sle1 values were normalized to the *cxaR*$_{down}$ band at 37 °C. The intensities, shown below the corresponding bands, represent the mean of two biological replicates, with standard deviation. (**B**) The effect of sublethal concentrations of penicillin G (1 ng/mL) on the cell wall-associated and extracellular levels of Sle1 in the NCTC8325-4 wild-type and Δ*cxaR* mutant, shown by Western blotting. Band intensities were quantified using ImageJ. Values for cell wall-associated Sle1 were normalized to the wild-type band without penicillin G, while extracellular Sle1 values were normalized to the Δ*cxaR* band without penicillin G. The intensities, shown below the corresponding bands, represent the mean of two biological replicates, with standard deviation. (**C**) Western blotting of cell wall-associated Sle1 in the NCTC8325-4 Δ*cxaR* mutant in response to sublethal concentrations of various antibiotics. From left to right; - (without antibiotics), OXA (7 ng/mL oxacillin), TEC (62.5 ng/mL teicoplanin), BAC (1 µg/mL bacitracin), MOE (7.5 ng/mL moenomycin), TUN (200 ng/mL tunicamycin), NOV (15 ng/mL novobiocin), KAN (300 ng/mL kanamycin). Band intensities were quantified using ImageJ. Values for cell wall-associated Sle1 were normalized to the Δ*cxaR* band without antibiotics. The intensities, shown below the corresponding bands, represent the mean of two biological replicates, with standard deviation.

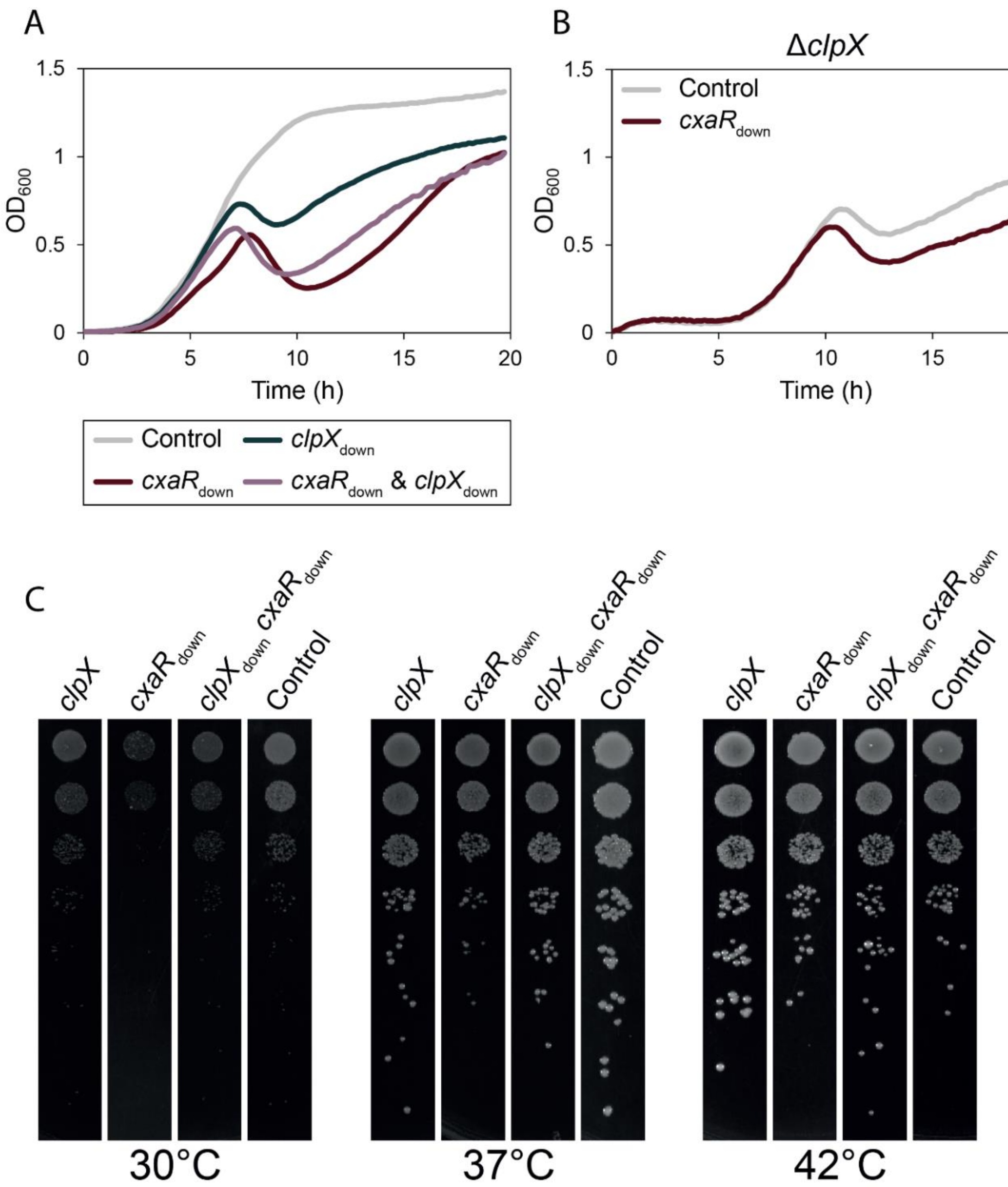


**S11 Fig. *cxaR* mutant cells resemble *clpX* mutants, exhibiting a comparable temperature-dependent growth defect.** (**A**) The effect of concurrent CRISPRi knockdown of *clpX* and *cxaR* (purple) in NCTC8325-4 compared to the CRISPRi control (grey) and single knockdown of *clpX* (green) or *cxaR* (red) during growth in TSB at 37°C. Gene knockdown was induced with addition of 500 µM IPTG. The graphs represent averages from triplicate measurements. (**B**) Growth of an NCTC8325-4 Δ*clpX* mutant in TSB at 37°C with (red) and without (grey) knockdown of *cxaR*. Gene knockdown was induced with addition of 500 µM IPTG. The graphs represent averages from triplicate measurements. (**C**) Spotting assay of cells with single knockdown of either *clpX* ($clpX_{down}$) or *cxaR* ($cxaR_{down}$), and double knockdown of both *clpX* and *cxaR* ($clpX_{down}$ $cxaR_{down}$) compared to the CRISPRi control at 30°C, 37°C, and 42°C. The strains were grown to mid-exponential phase ($OD_{600}$ of 0.5), 10-fold serially diluted, and spotted in volumes of 5 µL on TSA with 500 µM IPTG.

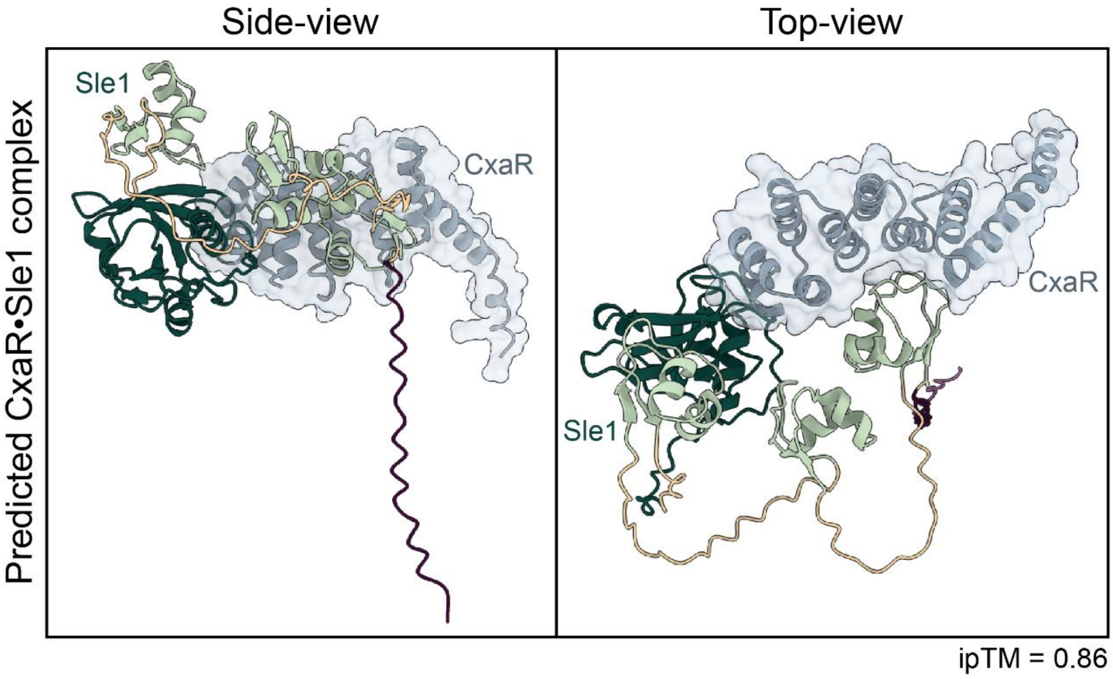


**S12 Fig. The AlphaFold 3-predicted interaction between CxaR and Sle1.** The domains of Sle1 are colored in accordance with the structure in **Fig. 6B**, with the signal peptide in purple, LysM domains in light green, and CHAP domain in dark green. The predicted complex has a high confidence interaction score (ipTM) of 0.86.

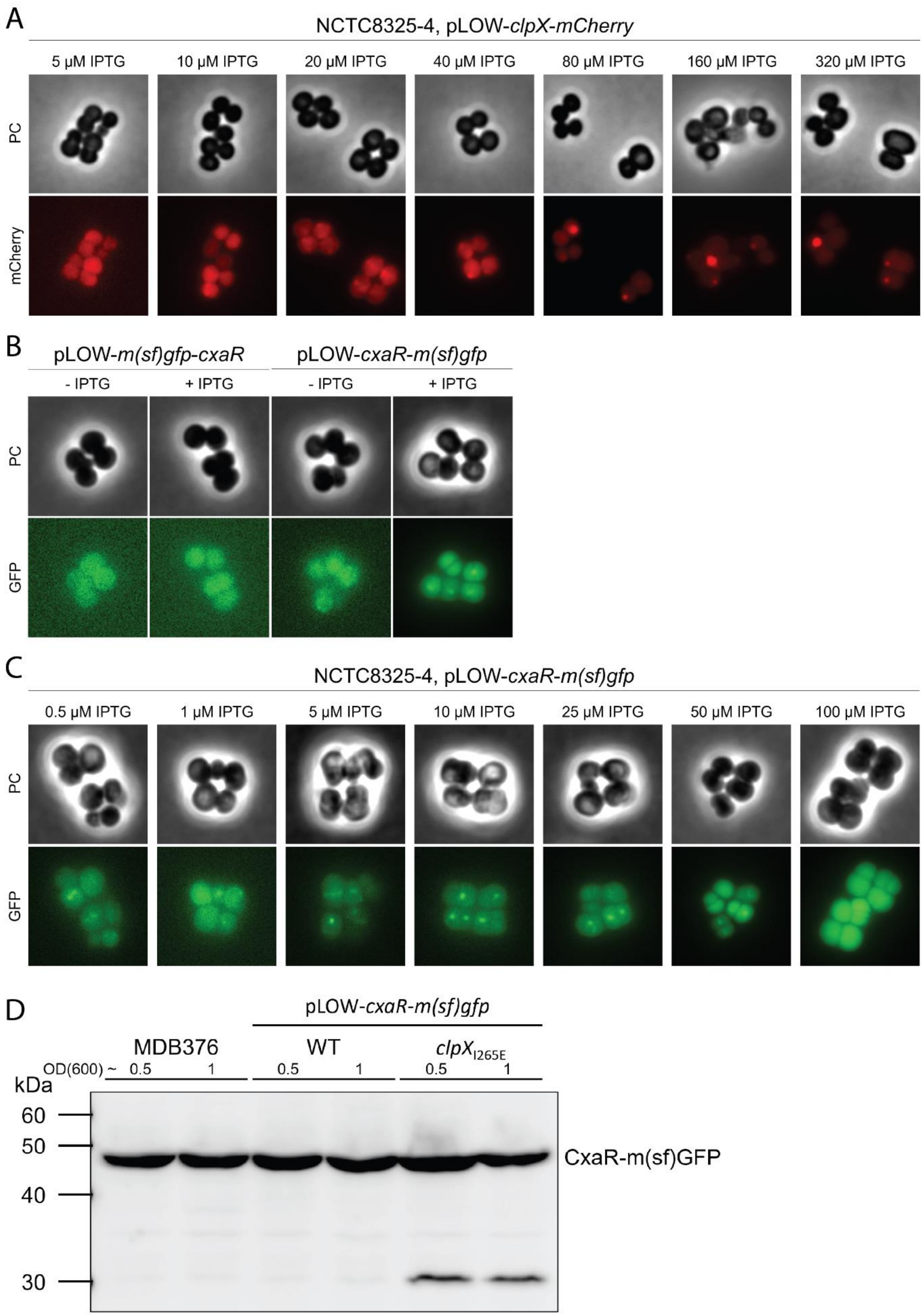


**S13 Fig. Fluorescent fusions of ClpX and CxaR.** (**A**) Phase contrast and fluorescence microscopy of a plasmid-borne C-terminal ClpX-mCherry fusion in NCTC8325-4, with inducer concentrations between 5 µM and 320 µM IPTG. (**B**) Phase contrast and fluorescence microscopy of NCTC8325-4 carrying plasmids encoding an N-terminal (left) and C-terminal (right) CxaR-GFP fusion, with and without induction of expression by 50 µM IPTG. (**C**) Various induction levels of the plasmid-borne C-terminal CxaR-GFP fusion in NCTC8325-4. The concentrations of the IPTG inducer ranged from 0.5 µM to 100 µM, and localization was assessed with phase contrast and fluorescence microscopy. (**D**) Western blot of the C-terminal CxaR-GFP fusion from whole cell lysate expressed from the native *cxaR* chromosomal locus in NCTC8325-4 (MDB376), from the pLOW plasmid in the NCTC8325-4 wild-type and from the pLOW plasmid in the NCTC8325-4 $clpX_{I265E}$ mutant. Expression of the fusion from pLOW was induced by 10 µM IPTG. Cultures were harvested at $OD_{600}$ 0.5 and 1.0.

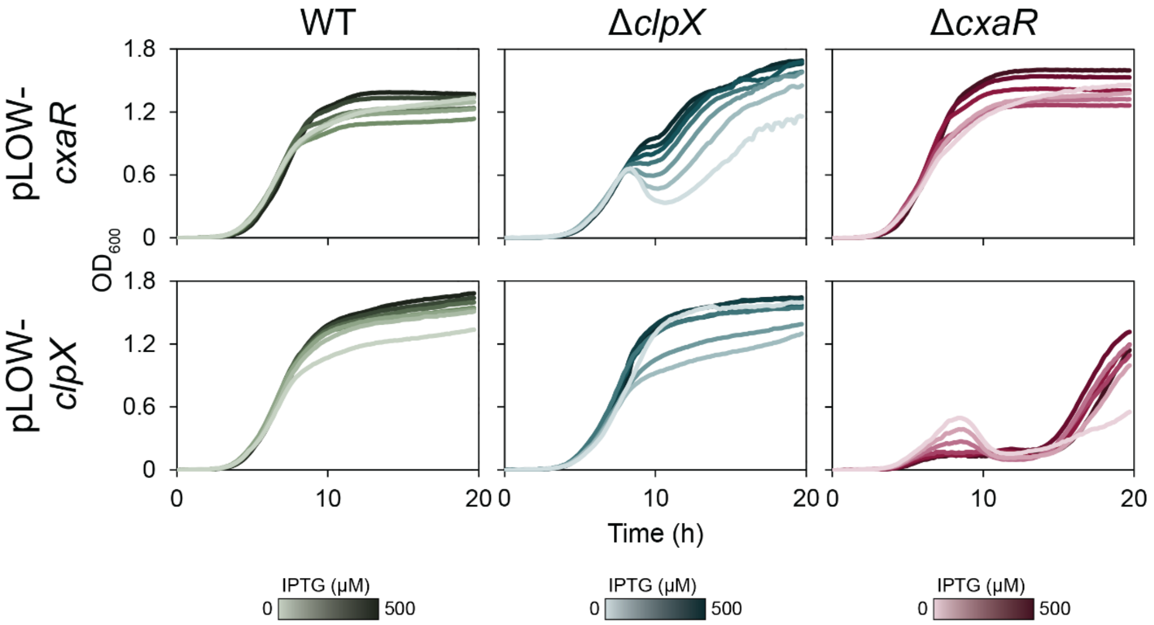


**S14 Fig. Overexpression of CxaR can compensate for the absence of ClpX, whereas ClpX overexpression further exacerbates the growth defect of Δ*cxaR*.** Overexpression of *cxaR* and *clpX* in the NCTC8325-4 wild-type (green), Δ*clpX* mutant (blue) and Δ*cxaR* mutant (red) from pLOW-*cxaR* and pLOW-*clpX*, respectively. Concentrations of IPTG ranged from 0 to 500 µM.

### Supporting Tables

**S1 Table. CRISPRi-seq identified genetic factors modulating susceptibility to β-lactams in *S. aureus*.** The CRISPRi library, targeting over 97% of the genomic features of *S. aureus* NCTC8325-4 [1], was transformed into a NCTC8325-4 strain where *dCas9* expression is controlled by a tetracycline-inducible promoter on the chromosome [2]. The induced library was grown in the presence and absence of 0.008 µg/ml penicillin G for ~20 generations at 37ºC in milk. Plasmids were subsequently isolated, and the sgRNA abundances were determined by Illumina sequencing, as previously described [3]. $Log_2$ fold-changes ($Log_2FC$) between penicillin G treated (PenG) and non-treated (minPenG) samples are presented for each sgRNA, along with the corresponding p-values ($P_{adj}$). Explanation of columns headings: sgRNA; sgRNA identification number, Locus tag; locus tag of the gene in *S. aureus* NCTC8325-4 targeted by the sgRNA, name; gene name, minPenG essentiality; whether the targeted gene is classified as essential, neutral or costly for growth without penicillin G added, PenG essentiality; whether the targeted gene is classified as essential, neutral or costly for growth with 0.008 µg/ml penicillin G added, Log2FC; log2 fold-change in sgRNA abundance between conditions with or without penicillin G; Padj; adjusted P-value, Interactions; whether (yes or no) there were significant (Padj < 0.05) difference in log2FC between conditions with or without penicillin G. Transcriptional units (sgRNAs) whose knockdown were found to significantly alter sensitivity to penicillin are labelled with colors, where blue indicates increased sensitivity and red indicates reduced sensitivity. *SAOUHSC_00659* (*cxaR*) is shown in bold.

**S2 Table. The spontaneous mutations acquired when the *cxaR* deletion mutant was grown at a 30°C.** List of mutations, with over 90% variant frequency, identified when re-sequencing five independently isolated, visibly larger NCTC8325-4 Δ*cxaR* colonies formed at 30°C.

| Strain name | Gene affected | Nucleotide change | Polymorphism type | Amino acid change | Protein effect | Variant Frequency |
|---|---|---|---|---|---|---|
| EFS104 | *aldA* | G>T | SNP (transversion) | E375D | Substitution | 99.5 % |
| | (RBS of *sle1*) | G>A (at position −13 relative to the *sle1* start codon) | SNP (transition) | | | 99.3 % |
| EFS105 | *sle1* | G>A | SNP (transition) | M292I | Substitution | 99.7 % |
| MDB390 | *sle1* | G>T | SNP (transversion) | G315C | Substitution | 100.0% |
| MDB401 | *sle1* | C>T | SNP (transition) | Q197* | Truncation | 99.7% |
| | *sarX* | C>T | SNP (transition) | H65Y | Substitution | 100.0% |
| MDB404 | *sle1* | (A)8>(A)7 | Deletion (tandem repeat) | V5* | Frame Shift | 99.7% |

**S3 Table.** Strains and mutants used in this work.

| Name | Genotype and characteristics[a] | Reference |
|---|---|---|
| ***S. aureus* NCTC8325-4** | | |
| NCTC8325-4/ MDB1 | MSSA lab strain, derivative of NCTC8325 cured of prophages | [4] |
| MM267 | NCTC8325-4, *tetR*_$P_{tet}$_*dCas9*, *tetM* | Lab collection |
| MHU16 | MM267 carrying pVL2336-sgRNA(*cxaR*), $cam^r$ | This work |
| MHU46 | MM267 carrying pVL2336-sgRNA(*S245*), $cam^r$ | This work |
| MHU48 | MM267 carrying pVL2336-sgRNA(*SAOUHSC_00660*), $cam^r$ | This work |
| MHU57 | MM267 carrying pVL2336-sgRNA(*SAOUHSC_00655*), $cam^r$ | This work |
| MHU58 | MM267 carrying pVL2336-sgRNA(*SAOUHSC_00656*), $cam^r$ | This work |
| MHU59 | MM267 carrying pVL2336-sgRNA(*SAOUHSC_00658*), $cam^r$ | This work |
| MK2047 | NCTC8325-4, Δ*cxaR*, $spc^r$ | This work |
| EFS104 | MK2047, *sle1* $RBS_{G\rightarrow A}$, $spc^r$ | This work |
| EFS105 | MK2047, $sle1_{M292I}$, $spc^r$ | This work |
| MDB390 | MK2047, $sle1_{G315C}$, $spc^r$ | This work |
| MDB401 | MK2047, $sle1_{Q197*}$, $spc^r$ | This work |
| MDB404 | MK2047, $sle1_{V5*}$, $spc^r$ | This work |
| MK2061 | MK2047 carrying pLOW-*dCas9*, $spc^r$, $ery^r$ | This work |
| MK2063 | MK2061 carrying pVL2236-sgRNA(*sle1*), $spc^r$, $ery^r$, $cam^r$ | This work |
| MK2066 | MK2061 carrying pCG248-sgRNA(*luc*), $spc^r$, $ery^r$, $cam^r$ | This work |
| MDB432 | MK2047 carrying pLOW-*cxaR*, $spc^r$, $ery^r$ | This work |
| MDB433 | MK2047 carrying pLOW-*cxaR-m(sf)gfp*, $spc^r$, $ery^r$ | This work |
| MDB447 | MK2047 carrying pLOW-*clpX-mCherry*, $spc^r$, $ery^r$ | This work |
| MDB429 | NCTC8325-4 carrying pLOW-*cxaR*, $ery^r$ | This work |
| MDB430 | NCTC8325-4 carrying pLOW-*cxaR-m(sf)gfp*, $ery^r$ | This work |
| MDB431 | NCTC8325-4 carrying pLOW-*m(sf)gfp-cxaR*, $ery^r$ | This work |
| MDB445 | NCTC8325-4 carrying pLOW-*clpX*, $ery^r$ | This work |
| MDB454 | NCTC8325-4 carrying pLOW-*clpX-mCherry*, $ery^r$ | This work |
| MDB376 | *cxaR-m(sf)gfp* (chromosomal), $spc^r$ | This work |
| MDB453 | MDB376 carrying pLOW-*clpX-mCherry*, $spc^r$, $ery^r$ | This work |
| MDB326 | NCTC8325-4, Δ*clpX* | [5] |
| EFS166 | MDB326 carrying pLOW-*dCas9*, $ery^r$ | This work |
| EFS168 | EFS166 carrying pVL2336-sgRNA(*cxaR*), $ery^r$, $cam^r$ | This work |
| EFS169 | EFS166 carrying pVL2336-sgRNA(*luc*), $ery^r$, $cam^r$ | This work |
| MDB394 | MDB326 carrying pLOW-*cxaR-m(sf)gfp*, $ery^r$ | This work |
| MDB435 | MDB326 carrying pLOW-*cxaR*, $ery^r$ | This work |
| MDB446 | MDB326 carrying pLOW-*clpX*, $ery^r$ | This work |
| MDB327 | NCTC8325-4, $clpX_{I265E}$ | [6] |
| EFS167 | MDB327 carrying pLOW-*dCas9*, $ery^r$ | This work |
| EFS170 | EFS167 carrying pVL2336-sgRNA(*cxaR*), $ery^r$, $cam^r$ | This work |
| EFS171 | EFS167 carrying pVL2336-sgRNA(*luc*), $ery^r$, $cam^r$ | This work |
| MDB437 | MDB327 carrying pLOW-*cxaR*, $ery^r$ | This work |
| MDB395 | MDB327 carrying pLOW-*cxaR-m(sf)gfp*, $ery^r$ | This work |
| MK1465 | NCTC8325-4 carrying pLOW-*dCas9*, $ery^r$ | Lab collection |
| EFS82 | MK1465 carrying pVL2336-sgRNA(*clpX*), $ery^r$, $cam^r$ | This work |
| EFS84 | MK1465 carrying pVL2336-sgRNA(*cxaR*), $ery^r$, $cam^r$ | This work |
| EFS68 | MK1465 carrying pVL2336-sgRNA(*cxaR+clpX*), $ery^r$, $cam^r$ | This work |
| EFS88 | MK1465 carrying pCG248-sgRNA(*luc*), $ery^r$, $cam^r$ | This work |

| | | |
|---|---|---|
| EFS141 | NCTC8325-4 carrying pAF256-*cxaR-SmBit/LgBit*, cam$^r$ | This work |
| EFS145 | NCTC8325-4 carrying pAP118-*cxaR-SmBit/clpP-LgBit*, cam$^r$ | This work |
| EFS148 | NCTC8325-4 carrying pAP118-*cxaR-SmBit/clpX-LgBit*, cam$^r$ | This work |
| EFS177 | NCTC8325-4 carrying pAP118-*cxaR-SmBit/cxaR-LgBit*, cam$^r$ | This work |
| MDB384 | NCTC8325-4 carrying pAP118-*cxaR-SmBit/sle1-LgBit*, cam$^r$ | This work |
| MDB385 | NCTC8325-4 carrying pAP118-*cxaR-SmBit/sle1$_{(no\ signal\ peptide)}$-LgBit*, cam$^r$ | This work |
| MDB417 | NCTC8325-4 carrying pAF256-*clpX-SmBit/LgBit*, cam$^r$ | This work |
| MDB418 | NCTC8325-4 carrying pAP118-*clpX-SmBit/sle1-LgBit*, cam$^r$ | This work |
| MDB419 | NCTC8325-4 carrying pAP118-*clpX-SmBit/sle1$_{(no\ signal\ peptide)}$-LgBit*, cam$^r$ | This work |
| MDB420 | NCTC8325-4 carrying pAP118-*clpX-SmBit/cxaR-LgBit*, cam$^r$ | This work |
| MDB421 | NCTC8325-4 carrying pAP118-*clpX-SmBit/clpP-LgBit*, cam$^r$ | This work |
| MDB466 | NCTC8325-4 carrying pAF256-*clpX$_{I265E}$-SmBit/LgBit*, cam$^r$ | This work |
| MDB469 | NCTC8325-4 carrying pAP118-*clpX$_{I265E}$-SmBit/cxaR-LgBit*, cam$^r$ | This work |
| MDB470 | NCTC8325-4 carrying pAP118-*clpX$_{I265E}$-SmBit/clpP-LgBit*, cam$^r$ | This work |
| ***S. aureus* JE2** | | |
| JE2/MDB9 | Community acquired MRSA strain, derivative of USA300 LAC cured of plasmids | [7] |
| MDB16 | JE2 carrying pLOW-*dCas9_aad9*, spc$^r$ | Lab collection |
| MHU44 | MDB16 carrying pVL2336-sgRNA(*cxaR*), spc$^r$, cam$^r$ | This work |
| MDB44 | MDB16 carrying pCG248-sgRNA(*luc*), spc$^r$, cam$^r$ | Lab collection |
| ***E. coli*** | | |
| IM08B | DH10B, Δ*dcm*, $P_{help}$-*hsdMS*, $P_{N25}$-*hsdS* (strain expressing the *S. aureus* CC8 specific methylation genes) | [8] |

**S4 Table.** Plasmids used in this work.

| Name | Description[a] | Reference |
|---|---|---|
| pLOW | Low-copy number staphylococcal shuttle vector with a IPTG inducible P*spac* promoter and *lacI* repressor, amp$^r$, ery$^r$ | [9] |
| pLOW-*dCas9_aad9* | For IPTG-inducible expression of dCas9, amp$^r$, spc$^r$ | [10] |
| pLOW-*dCas9*_extra_*lacO* | For IPTG-inducible expression of dCas9, amp$^r$, ery$^r$ | [11] |
| pLOW-*cxaR* | For IPTG-inducible expression of CxaR, amp$^r$, ery$^r$ | This work |
| pLOW-*m(sf)gfp-cxaR* | For IPTG-inducible expression of CxaR with GFP fused to its N-terminus, amp$^r$, ery$^r$ | This work |
| pLOW-*cxaR-m(sf)gfp* | For IPTG-inducible expression of CxaR with GFP fused to its C-terminus, amp$^r$, ery$^r$ | This work |
| pLOW-*clpX* | For IPTG-inducible expression of CxaR, amp$^r$, ery$^r$ | This work |
| pLOW-*clpX-mCherry* | For IPTG-inducible expression of CxaR with mCherry fused to its C-terminus, amp$^r$, ery$^r$ | This work |
| pCG248 | *E. coli*/*S. aureus* shuttle vector, amp$^r$, cam$^r$ | [12] |
| pCG248-sgRNA(*luc*) | For constitutive expression of a nontargeting sgRNA, amp$^r$, cam$^r$ | [11] |
| pVL2336 | *E. coli*/*S. aureus* shuttle vector, amp$^r$, cam$^r$ | [13] |
| pVL2336-sgRNA(*cxaR*) | For constitutive expression of sgRNA targeting *cxaR*, amp$^r$, cam$^r$ | This work |
| pVL2336-sgRNA(*clpX*) | For constitutive expression of sgRNA targeting *clpX*, amp$^r$, cam$^r$ | This work |
| pVL2336-sgRNA(*cxaR*+*clpX*) | For constitutive expression of sgRNA targeting both *cxaR* and *clpX*, amp$^r$, cam$^r$ | This work |
| pVL2336-sgRNA(*SAOUHSC_00655*) | For constitutive expression of sgRNA targeting *SAOUHSC_00655*, amp$^r$, cam$^r$ | This work |
| pVL2336-sgRNA(*SAOUHSC_00656*) | For constitutive expression of sgRNA targeting *SAOUHSC_00656*, amp$^r$, cam$^r$ | This work |
| pVL2336-sgRNA(*SAOUHSC_00658*) | For constitutive expression of sgRNA targeting *SAOUHSC_00658*, amp$^r$, cam$^r$ | This work |
| pVL2336-sgRNA(*SAOUHSC_00660*) | For constitutive expression of sgRNA targeting *SAOUHSC_00660*, amp$^r$, cam$^r$ | This work |
| pVL2336-sgRNA(*S245*) | For constitutive expression of sgRNA targeting *S245*, amp$^r$, cam$^r$ | This work |
| pVL2336-sgRNA(*sle1*) | For constitutive expression of sgRNA targeting *sle1*, amp$^r$, cam$^r$ | This work |
| pMAD | Thermosensitive shuttle vector for allelic replacement in Gram-positive bacteria, amp$^r$, ery$^r$ | [14] |
| pMAD-*cxaR::spc* | To knock out *cxaR* by allelic replacement, amp$^r$, ery$^r$ | This work |
| pMAD-*cxaR-m(sf)gfp_spc* | To chromosomally integrate GFP-tagged *cxaR* in its native locus, amp$^r$, ery$^r$ | This work |
| pAF256-P$_{tet}$-*hupA-smbit/lgbit* | Split-luciferase control plasmid, originally for assessing HupA self-interaction in *C. difficile*, cam$^r$ | [15] |
| pAF256-*cxaR-smbit/lgbit* | CxaR split-luciferase control, cam$^r$ | This work |
| pAF256-*clpX-smbit/lgbit* | ClpX split-luciferase control, cam$^r$ | This work |
| pAF256-*clpX$_{I265E}$-smbit/lgbit* | ClpX$_{I265E}$ split-luciferase control, cam$^r$ | This work |
| pAP118-P$_{tet}$-*hupA-smbit/hupA-lgbit* | Split-luciferase test plasmid, originally for assessing HupA self-interaction in *C. difficile*, cam$^r$ | [15] |
| pAP118-*cxaR-smbit/clpP-lgbit* | For assessing interactions between CxaR and ClpP, cam$^r$ | This work |
| pAP118-*cxaR-smbit/clpX-lgbit* | For assessing interactions between CxaR and ClpX, cam$^r$ | This work |
| pAP118-*cxaR-smbit/cxaR-lgbit* | For assessing CxaR self-interaction, cam$^r$ | This work |
| pAP118-*cxaR-smbit/sle1-lgbit* | For assessing interactions between CxaR and Sle1, cam$^r$ | This work |

| | | |
|---|---|---|
| pAP118-*cxaR-smbit/sle1*$_{(no\ signal\ peptide)}$-*lgbit* | For assessing interactions between CxaR and Sle1, cam$^r$ | This work |
| pAP118-*clpX-smbit/sle1-lgbit* | For assessing interactions between ClpX and Sle1, cam$^r$ | This work |
| pAP118-*clpX-smbit/sle1*$_{(no\ signal\ peptide)}$-*lgbit* | For assessing interactions between ClpX and Sle1, cam$^r$ | This work |
| pAP118-*clpX-smbit/cxaR-lgbit* | For assessing interactions between ClpX and CxaR, cam$^r$ | This work |
| pAP118-*clpX-smbit/clpP-lgbit* | For assessing interactions between ClpX and ClpP, cam$^r$ | This work |
| pAP118-*clpX*$_{I265E}$-*smbit/cxaR-lgbit* | For assessing interactions between ClpX$_{I265E}$ and CxaR, cam$^r$ | This work |
| pAP118-*clpX*$_{I265E}$-*smbit/clpP-lgbit* | For assessing interactions between ClpX$_{I265E}$ and ClpP, cam$^r$ | This work |

a. amp$^r$ = ampicillin resistant, ery$^r$ = erythromycin resistant, spc$^r$ = spectinomycin resistant, cam$^r$ = chloramphenicol resistant.

**S5 Table.** Oligonucleotide primers used in this work.

| Name | Sequence 5'-3'[a] | Description[b] |
|---|---|---|
| **Primers for construction of pMAD-*cxaR::spc*** | | |
| mk188 | ATTGGGCCCACCTAGGATC | *aad9*_up F |
| mk189 | ACTATGCGGCCGCTCGAG | *aad9*_down R |
| mk591 | ATCGCCATGGGCAGATAATACAAAAGATATGGT | *cxaR*_up F w/NcoI |
| mk592 | **GATCCTAGGTGGGCCCAAT**CGTACTAAATCAAACTCGCCA | *cxaR*_up R |
| mk593 | **CTCGAGCGGCCGCATAGT**ATTACACCAACGTAATGCACG | *cxaR*_down F |
| mk594 | ATCGGGATCCTCGCTTCAAAGGCAGTACCA | *cxaR*_down R w/BamHI |
| **Primers for construction of pMAD-*cxaR-m(sf)gfp_spc*** | | |
| mdb58 | ACCTCCATGGGATCCAGGTGCTCAAAGTATG | *cxaR*_up F w/NcoI |
| mdb59 | ATGTTTTTTTATAAGGTGTTGCAT | *cxaR*_up R |
| mdb60 | ATGCAACACCTTATAAAAAAACAT | *cxaR-m(sf)gfp* F |
| mdb61 | **TTTCCGTTAATCAAATTGCTCAT**TTACTTATAAAGCTCATCCATGC | *cxaR-m(sf)gfp* R |
| mdb62 | **CTATAGAAACTTCTCTCAATTAG**TTACATTTCAATTATATTAGCTTAAT | *cxaR*_down F |
| mdb63 | AGGTGTCGACCCAAACCAGATTCCCTTCCA | *cxaR*_down R w/SalI |
| mk503 | ATGAGCAATTTGATTAACGGAAA | *spc* F |
| mk504 | CTAATTGAGAGAAGTTTCTATAG | *spc* R |
| **Primers for screening of pMAD plasmids** | | |
| im156 | AATCTAGCTAATGTTACGTTACA | pMAD F |
| mk177 | GATGCCGCCGGAAGCGAG | pMAD R |
| **Primers for creating pLOW constructs** | | |
| mhu26 | ACGTGGATCCCAACACCTTATAAAAAAACATGTA | *cxaR* F w/BamHI |
| mhu27 | ACGTGAATTCATGTAATTAACGTGCATTACGTTG | *cxaR* R w/EcoRI |
| mhu28 | ATCCTCGAGTCGACCAATAAAACT*AGGAGG*AAATTTAAATGCAACAC CTTATAAAAAAACAT | *cxaR* F w/XhoI+SalI |
| mhu29 | TCCGGGGATCCGACGTCCATTACGTTGGTGTA | *cxaR* R w/BamHI |
| mdb98 | ATCCTCGAGGTCGACCAATAAAACT*AGGAGG*AAATTTAAATGTTTAA ATTCAATGAAGATGAA | *clpX* F w/XhoI+SalI |
| mdb99 | ACGTGAATTCCAATGATTAAGCTGATGTTTTAC | *clpX* R w/EcoRI |
| mdb102 | TCCGGGGATCCAAGCTGATGTTTTACTATTATTAATT | *clpX* R w/BamHI |
| **Primers for screening of pLOW plasmids** | | |
| im218 | TCTCATTCAATTCCTAGGTGG | pLOW F |
| im134 | TGTGCTGCAAGGCGATTAAG | pLOW R |
| **Oligos for creating pVL2336-sgRNA constructs** | | |
| SAOUHSC_00427_for | TATAATTGCCCACACTGATTCACC | *sle1* sgRNA F |
| SAOUHSC_00427_rev | AAACGGTGAATCAGTGTGGGCAAT | *sle1* sgRNA R |
| SAOUHSC_00655_for | TATATAGCAATCAGATCTAACTCT | *_00655* sgRNA F |
| SAOUHSC_00655_rev | AAACAGAGTTAGATCTGATTGCTA | *_00655* sgRNA R |
| SAOUHSC_00656_for | TATACACGAACCATGTTAACCCCG | *_00656* sgRNA F |
| SAOUHSC_00656_rev | AAACCGGGGTTAACATGGTTCGTG | *_00656* sgRNA R |
| SAOUHSC_00658_for | TATACACTTGCAATTTCTTTACTG | *_00658* sgRNA F |
| SAOUHSC_00658_rev | AAACCAGTAAAGAAATTGCAAGTG | *_00658* sgRNA R |
| SAOUHSC_00659_for | TATAAAATTCCATAAAATCTGTTT | *cxaR* sgRNA F |
| SAOUHSC_00659_rev | AAACAAACAGATTTTATGGAATTT | *cxaR* sgRNA R |

| | | |
|---|---|---|
| SAOUHSC_ S245_for | TATATTTTAAGTATTAAAAAAGGC | *S245* sgRNA F |
| SAOUHSC_ S245_rev | AAACGCCTTTTTTAATACTTAAAA | *S245* sgRNA R |
| SAOUHSC_ 00660_for | TATAGCCCACTACAATGCCGACGT | _*00660* sgRNA F |
| SAOUHSC_ 00660_rev | AAACACGTCGGCATTGTAGTGGGC | _*00660* sgRNA R |
| SAOUHSC_ 01778_for | TATATCCATAATTTCTTTAGGAGT | *clpX* sgRNA F |
| SAOUHSC_ 01778_rev | AAACACTCCTAAAGAAATTATGGA | *clpX* sgRNA R |
| **Primers for screening of pCG248 and pVL2336 plasmids** | | |
| mk26 | GGATAACCGTATTACCGCCT | pCG248 F |
| mk25 | AAATCTCGAAAATAATAGAGGGA | pCG248 R |
| **Primers for creating split-luciferase constructs** | | |
| efs7 | TCAGAGCTCATCTGGAGAAAT*AGGAGG*AC | *cxaR* F w/SacI |
| efs8 | GCCCTCGAGAACGTGCATTACGTTGGTGTA | *cxaR* R w/XhoI |
| mdb93 | ATCGGAGCTCCAATCTAGTATAGTCTTTAACG | *clpX* F w/SacI |
| mdb94 | GCCCTCGAGCAGCTGATGTTTTACTATTATTAAT | *clpX* R w/XhoI |
| efs5 | GTCCGATCGCAATCTAGTATAGTCTTTAACG | *clpX* F w/PvuI |
| efs6 | GGCGGCCGCAGCTGATGTTTTACTATTATTAAT | *clpX* R w/NotI |
| efs9 | GTCCGATCGGTAACAGTTATTACA*AGGAGG* | *clpP* F w/PvuI |
| efs10 | GGCGGCCGCTTTTGTTTCAGGTACCATCAC | *clpP* R w/NotI |
| efs20 | GTCCGATCGATCTGGAGAAAT*AGGAGG*AC | *cxaR* F w/PvuI |
| efs21 | GGCGGCCGCACGTGCATTACGTTGGTGTA | *cxaR* R w/NotI |
| mdb88 | ATCGCGATCGGTTAAGCAAG*AGGAGG*ATTTTA | *sle1* F w/PvuI |
| mdb89 | GCCGCGGCCGCGTGAATATATCTATAATTATTTACTT | *sle1* R w/NotI |
| mdb90 | ATCGCGATCGCAATAAAACTA*GGAGG*AAATTTAAATGGCTACAACTC ACACAGTAAAAC | *sle1* F w/PvuI |
| **Primers for screening of split-luciferase constructs** | | |
| efs11 | CATGCCAATACAATGTAGGC | Split-luciferase F |
| efs53 | AGACTTCTCATGAGAGAAGC | Split-luciferase R |

a. The restriction sites are underlined, the overhangs are bolded, and the ribosomal binding sites are italicized.
b. F = forward primer, R = reverse primer, RS = restriction site, and RBS = ribosomal binding site.
